# Identifying candidate detoxification genes in the ecdysteroid kinase-like (EcKL) and cytochrome P450 gene families in *Drosophila melanogaster* by integrating evolutionary and transcriptomic data

**DOI:** 10.1101/2020.02.17.951962

**Authors:** Jack L. Scanlan, Rebecca S. Gledhill-Smith, Paul Battlay, Charles Robin

## Abstract

The capacity to detoxify toxic compounds is essential for adaptation to the ecological niches of many organisms, especially insects. However, detoxification in insects is often viewed through the lens of mammalian detoxification research, even though the organ and enzyme systems involved have diverged for over half a billion years. Phosphorylation is a non-canonical phase II detoxification reaction that, among animals, occurs near exclusively in insects, but the enzymes responsible have never been cloned or otherwise identified. We propose the hypothesis that members of the arthropod-specific ecdysteroid kinase-like (EcKL) gene family encode detoxicative kinases. To test this hypothesis, we annotated the EcKL gene family in 12 species of *Drosophila* and explored their evolution within the genus. Many ancestral EcKL clades are evolutionarily unstable and have experienced repeated gene gain and loss events, while others are conserved as single copy orthologs. Leveraging multiple published gene expression datasets from *D. melanogaster*, and using the cytochrome P450s—a canonical detoxification family—as a test case, we demonstrate relationships between xenobiotic induction, detoxification tissue-enriched expression and evolutionary instability in the EcKLs and the P450s. We also found previously unreported genomic and transcriptomic variation in a number of EcKLs and P450s associated with toxic stress phenotypes using a targeted phenome-wide association study (PheWAS) approach. Lastly, we devised a systematic method for identifying candidate detoxification genes in large gene families that is concordant with experimentally determined functions of P450 genes in *D. melanogaster*. Applying this method to the EcKLs suggested a significant proportion of these genes play roles in detoxification, and that the EcKLs may constitute a detoxification gene family in insects. Additionally, we estimate that between 11–16 uncharacterised *D. melanogaster* P450s are strong detoxification candidates.

**Highlights:**

- The poorly characterised ecdysteroid kinase-like (EcKL) gene family is hypothesised to encode enzymes responsible for detoxification by phosphorylation in insects.
- An integrative ‘detoxification score’ method accurately categorises the known functions of a canonical detoxification family, the cytochrome P450s, and suggests many EcKLs are also involved in detoxification.
- A targeted phenome-wide association study finds novel associations between EcKL/P450 variation and a number of toxic stress phenotypes, such as two unlinked EcKL paralogs that are both associated with developmental methylmercury resistance.

## 1. Introduction

Toxins play central roles in competition and trophic interactions between species— this is especially true for insects, where they often define ecological niches and can limit the food sources species can exploit. A well-known example is the wide diversity of toxins produced by plants, which aim to kill or otherwise dissuade herbivorous insects feeding on their tissues; the capacity of an insect to tolerate these toxins partially defines which plants it can consume (Mithöfer and Boland, 2012) and contributes to plant-insect co-evolution (Edger et al., 2015; Ehrlich and Raven, 1964). Toxin tolerance has many components, one of the most well-studied of which is metabolic detoxification. In animals, detoxification is carried out by collections of enzyme and transporter systems present in a number of organs and tissues throughout the body, which were originally conceptualised as a series of “phases” that result in the sequential modification (phase I), conjugation (phase II) and excretion (phase III) of toxins. This “canonical model” of detoxification, as well as the typical reactions expected in each phase, were defined in the context of mammalian pharmacological research (Williams, 1959), and have been adopted by researchers of other animal taxa, including insects (Berenbaum and Johnson, 2015; Chahine and O’Donnell, 2011; Yu, 2008). However, one concern about this canonical model is that a mammalian lens may limit our understanding of insect detoxification, as the lineages leading to arthropods and vertebrates diverged approximately 600 m.y.a. (Reis et al., 2015). A consequence of this may be differences in the enzyme families used between taxa, as many gene families are not conserved over deep evolutionary time (Danchin et al., 2006; Lespinet et al., 2002).

One such non-conserved reaction appears to be xenobiotic phosphorylation, which is rare in mammals, but is common in bacteria and insects (S. C. Mitchell, 2015; Ramirez and Tolmasky, 2010; Wilkinson, 1986). Phosphorylation has the potential to be a phase II detoxification reaction, as phosphate groups are highly polar and can be conjugated to hydroxyl moieties, and the formation of phosphorylated metabolites of xenobiotic compounds has been observed in at least 18 insect species across seven insect orders (Table 1). Many of these metabolites were discovered by John Smith and colleagues in the late 1960s to early 1970s (Binning et al., 1967; Darby et al., 1966; Heenan and Smith, 1974; Smith and Turbert, 1964) before the widespread adoption of modern analytical methods like LC-MS, but more recent papers have characterised phosphate conjugates in some detail. A major metabolic pathway for the drug terfenadine in the desert locust, *Schistocerca gregaria*, is the phosphorylation of hydroxylated metabolites (Olsen et al., 2014), and the same species also metabolises the drug midazolam to glucose-phosphate conjugates, seemingly by phosphorylating the glucose moiety of glucose conjugates (Olsen et al., 2015). In perhaps a more ecologically relevant example, caterpillars of the gypsy moth, *Lymantria dispar*, phosphorylate the glycoside moiety of salicinoids found the leaves of a host plant, *Populus tremula* x *tremuloides*, a hybrid poplar tree. These phosphate conjugates—formed in the gut and perhaps also the Malpighian tubules— comprise a substantial proportion of excreted salicinoid-like compounds, especially when caterpillars are previously fed poplar leaves, suggesting this detoxification process is induced by the presence of poplar secondary metabolites (Boeckler et al., 2016). Likewise, the detoxicative kinase activity identified in *Gromphadorhina portentosa*, the Madagascar cockroach, is present in the midgut, fat body and Malpighian tubules of the insect and is inducible by *in vivo* exposure to phenobarbital (Gil et al., 1974; R. S. H. Yang and Wilkinson, 1973), a compound commonly used to induce detoxification gene expression (Misra et al., 2011; Willoughby et al., 2006). Phytoecdysteroid detoxification may also involve phosphorylation (Rharrabe et al., 2007), but the phosphorylation of ingested ecdysteroids has been studied in only one species (Modde et al., 1984), even though ecdysteroids can be phosphorylated *in vitro* with midgut tissue homogenates from a handful of other species (Webb et al., 1996; 1995; Weirich et al., 1986). Overall, many insects appear able to phosphorylate xenobiotic phenols, glycosides and/or steroids directly, and it is possible other xenobiotics may be metabolised in a similarly direct manner, or after hydroxylation or hydrolysis. To date, however, no detoxicative phosphotransferase enzymes have been cloned or otherwise identified at the genetic level. In this paper, we wish to highlight this outstanding question in insect toxicology and raise and test a hypothesis about the identity of these unknown enzymes.

**Table 1:**
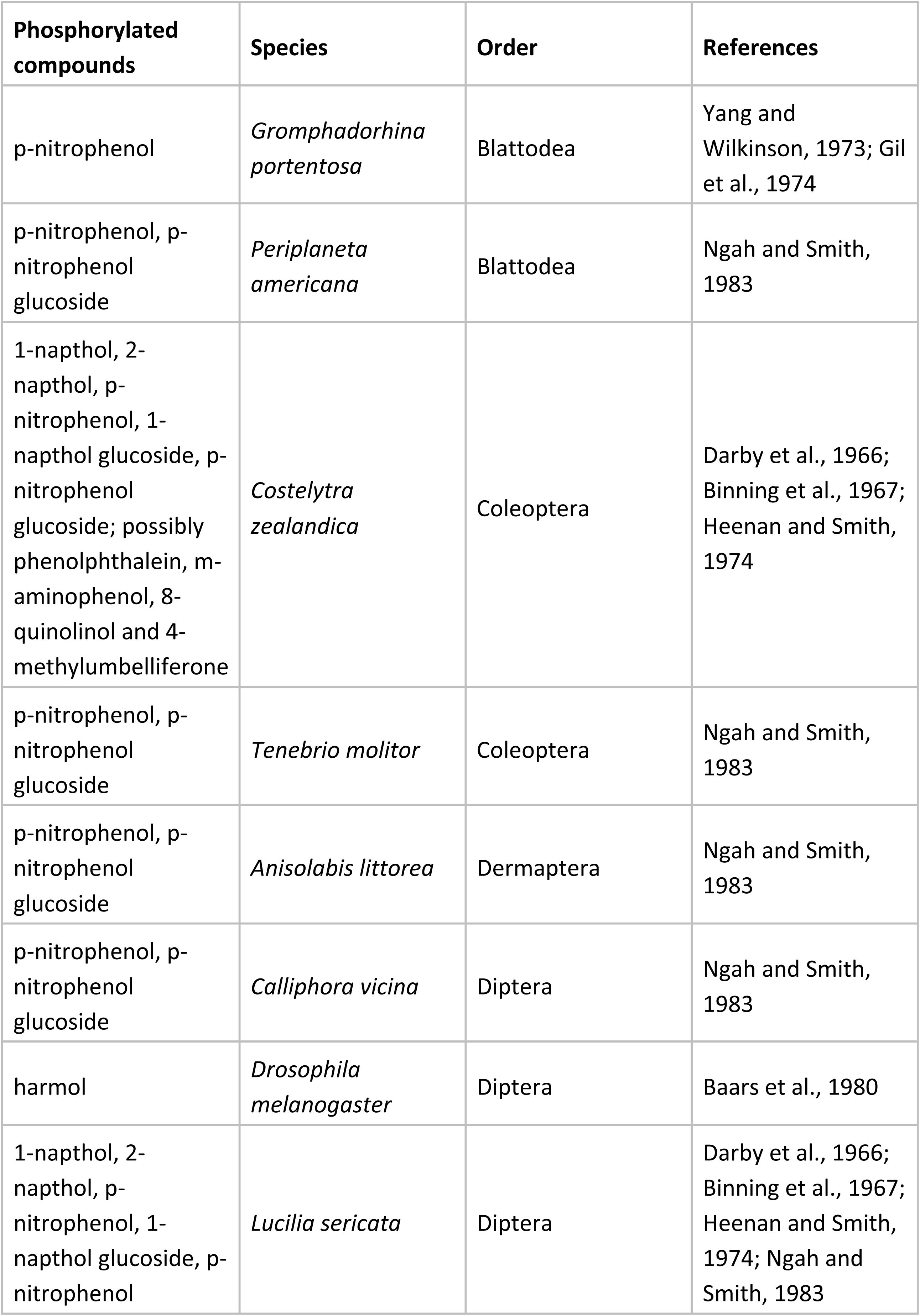
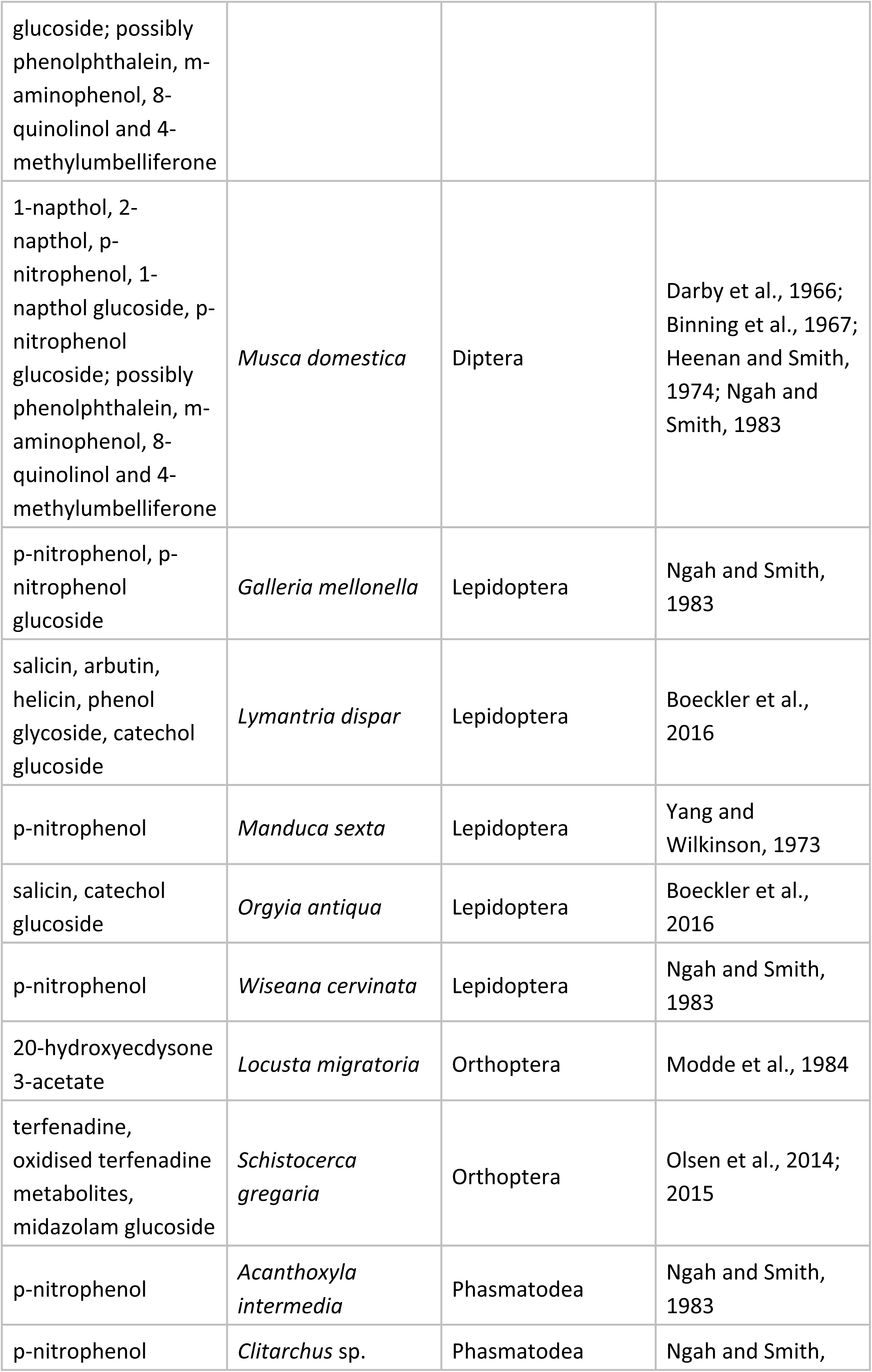
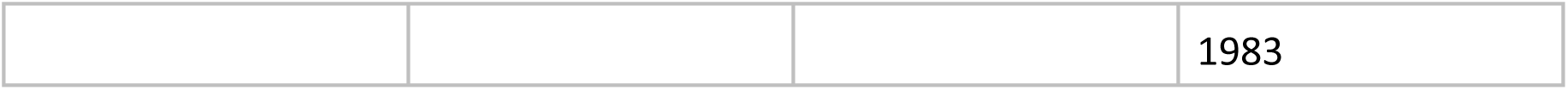
Phosphorylated metabolites of xenobiotic compounds in insects reported in the literature.

The identification of detoxification genes and enzymes is an important part of bridging the gap between toxicology, chemical ecology and functional genomics in insects. In the past, the main method of finding detoxification enzymes involved cloning and biochemical characterisation based on a known detoxification reaction. However, detoxification genes tend to have other characteristic properties, including transcriptional induction by xenobiotics (Willoughby et al., 2006) and enriched expression in tissues with known detoxification roles, like the midgut, Malpighian tubules and fat body (J. Yang et al., 2007). Phenotypes discovered during genetic experiments can also indicate detoxification functions, including susceptibility via gene disruption (H. Wang et al., 2018) or resistance via overexpression (Daborn et al., 2007); genome- and transcriptome-wide association studies can also identify candidate genes for detoxification-related phenotypes using naturally occurring variation (Robin et al., 2019). Additionally, genes encoding detoxification enzymes are thought to undergo gene duplication and loss at a faster rate than those encoding enzymes with important housekeeping functions (Kawashima and Satta, 2014; J. H. Thomas, 2007). The broad availability of “-omic” data in some insect taxa, particularly the *Drosophila* genus, raises the possibility that candidate detoxification genes could be identified by integrating evolutionary and transcriptomic data without any prior knowledge of biochemical functions. However, to our knowledge, this has yet to be attempted in a systematic way.

To validate this approach, we examine the cytochrome P450s (henceforth P450s), which are a large multigene family of enzymes that largely function as monooxygenases and catalyse phase I detoxification reactions such as hydroxylation, although a subset also catalyse a wider variety of reactions, including dealkylation, epoxidation and reduction (Bernhardt, 2006). P450s are an established detoxification family in virtually all animals, including insects (Heidel-Fischer and Vogel, 2015; Yu, 2008), and the number of P450 genes per genome in insects varies dramatically, from 38 in the fig wasp *Ceratosolen solmsi* to 222 in the little fire ant *Wasmannia auropunctata* (Rane et al., 2019). It has been suggested that diversity in the size of the P450 gene family may be linked to differences in detoxification capacity between taxa (Calla et al., 2017; Rane et al., 2019; 2016), although this has not been rigorously studied.

Multiple attempts have been made to classify P450s according to their biological functions. Thomas (2007) suggests a split between “endogenous-substrate” and “xenobiotic-substrate” enzymes, while Kawashima & Satta (2014) suggest a similar split with “biosynthesis-type” and “detoxification-type.” For the purposes of this paper, we adopt the classification system proposed by Gotoh (2012) of xenobiotic (X-class), secondary (S-class) and endogenous (E-class) functions. E-class enzymes synthesise or degrade compounds that are important for largely niche-independent developmental or physiological processes, such as hormones and cuticular hydrocarbons; S-class enzymes synthesise or degrade secondary metabolites that have niche-dependent functions not required for general viability, such as defensive compounds or pigments; and X-class enzymes detoxify xenobiotic compounds, which are also highly niche-dependent. E-and X-class P450s have been studied for many decades. A classic example of E-class enzymes are the Halloween P450s, which synthesise ecdysteroid moulting hormones from dietary sterols (Rewitz et al., 2007), but other E-class P450s belong to the biosynthetic pathways of juvenile hormones (Christesen et al., 2016; Helvig et al., 2004) and cuticular hydrocarbons (Qiu et al., 2012). Many X-class P450s have also been characterised and implicated in the detoxification of both natural and synthetic toxins (X. Li et al., 2007). S-class P450s are the least understood of the three classes, although some are involved in the biosynthesis of cyanogenic glycosides (Beran et al., 2019) and the degradation of pheromones (Wojtasek and Leal, 1999).

The evolutionary dynamics of P450s have been well-studied in insects and the *Drosophila* genus specifically (Feyereisen, 2011; 2006; Good et al., 2014). Within *Drosophila*, 30 ancestral P450 clades are stable—that is, they contain 1:1 orthologs in all studied species—while 30 clades have gene gain and gene loss in the genus, and 17 have only gene loss (Good et al., 2014). The genome of *Drosophila melanogaster* specifically contains 87 P450 genes, some of which have been studied in great detail (Chung et al., 2009). While evolutionary stability and developmentally essential (E-class) functions are thought to be linked in *D. melanogaster* (Chung et al., 2009), the link between evolutionary instability and X-class or S-class functions has yet to be rigorously established in this species.

The ecdysteroid kinase-like (EcKL; Interpro entry IPR004119) gene family is taxonomically restricted, being predominantly present in insect and crustacean genomes (A. Mitchell et al., 2014). EcKL enzymes are predicted to conjugate phosphate to secondary alcohols, using ATP as a phosphodonor (EC 2.7.1.-) and contain the EcKinase domain (Pfam accession PF02958), formerly known as DUF227 (domain of unknown function 227), a member of the Protein Kinase superfamily (CDD accession cl21453; El-Gebali et al., 2018). This superfamily contains protein kinases, as well as kinases with small molecular substrates, such as choline/ethanolamine kinases, aminoglycoside 3’-phosphotransferases and phosphoinositide 3-kinases (Marchler-Bauer et al., 2015). The EcKLs are currently named after a single member, *BmEc22K*, which encodes an ecdysteroid 22-kinase in the silkworm, *Bombyx mori*. BmEc22K phosphorylates the C-22 hydroxyl group of ecdysteroids, producing physiologically inactive ecdysteroid 22-phosphate conjugates (Sonobe et al., 2006). These conjugates are stored in the oocyte where they bind to vitellin in the yolk, and are hydrolysed to their active free form by an ecdysteroid-phosphate phosphatase (EPPase) after fertilisation to supply the ecdysteroid titre required for embryonic development (Sonobe and Yamada, 2004; Yamada and Sonobe, 2003; Yamada et al., 2005). This reciprocal conversion process may also occur in other insects and crustaceans, as orthopteran and other lepidopteran species also store ecdysteroid-phosphate conjugates in their eggs (Feldlaufer et al., 1987; Isaac and Rees, 1984; Isaac et al., 1983; Sonobe and Ito, 2009), as may some crustaceans (Subramoniam, 2000; Young et al., 1991). However, no ecdysteroid kinase/EPPase system has been molecularly characterised in any species besides *B. mori*, although EPPase orthologs exist in many insect genomes (Sonobe and Ito, 2009) and at least one crustacean genome (Asada et al., 2014).

No EcKLs besides BmEc22K have had their substrates identified, and very few other EcKLs have been functionally characterised. *Juvenile hormone-inducible protein 26* (*JhI-26*) is an EcKL found in *D. melanogaster* that has been implicated in *Wolbachia*-mediated cytoplasmic incompatibility (Liu et al., 2014). *CHKov1* and *CHKov2* are also EcKLs in *D. melanogaster*; a *CHKov1* allele containing a transposable element (TE) insertion confers resistance to the vertically-transmitted sigma virus, and a derived allele containing complete and partial duplications of both the *CHKov1*-TE allele and its neighbouring gene *CHKov2* confers an even greater level of resistance (Magwire et al., 2011). The mechanism underlying the resistance conferred by *CHKov1*-TE is currently unknown, although it is important to note that the TE insertion likely destroys the kinase function of the encoded polypeptides.

We raise the hypothesis that members of the EcKL gene family encode kinases responsible for the detoxicative phosphorylation seen in insects, based on four observations: first, the apparent taxonomic distribution of EcKLs is consistent with the limited taxonomic distribution of detoxicative phosphorylation in animals (S. C. Mitchell, 2015; Scanlan and Robin, unpublished); second, ecdysteroid kinase activity has been linked to detoxicative phosphorylation of phytoecdysteroids in some insects (Rharrabe et al., 2007); third, the size of the EcKL family appears to vary considerably between insect taxa (A. Mitchell et al., 2014; Scanlan and Robin, unpublished), suggesting not all members encode E-class enzymes; and fourth, EcKLs are at least distantly related to the aminoglycoside 3’-phosphotransferases, known detoxicative phosphotransferase enzymes (Marchler-Bauer et al., 2015).

Here we conduct the first evolutionary analysis of the EcKL gene family, focused on the dipteran genus *Drosophila*, demonstrating there is wide variability in the stability of EcKL orthologs within this taxon. We then show that integrating evolutionary, xenobiotic induction, transcriptional regulation and tissue expression datasets can be used to accurately predict which members of the P450 gene family are involved in xenobiotic detoxification, and apply this method to the EcKLs, demonstrating it is a strong candidate for a non-canonical part of the insect detoxification system. We also show, using a targeted phenome-wide association study (PheWAS) approach in the Drosophila Genetic Reference Panel (DGRP), that EcKL and P450 genomic and transcriptomic variation is associated with toxic stress phenotypes, providing candidate detoxification functions for members of these two gene families. Finally, we perform RNAi knockdown on a subset of EcKLs in *D. melanogaster* to find developmental lethality phenotypes and identify candidate E-class genes in this gene family.

## 2. Materials and methods

### 2.1. Gene family annotation

Eleven *Drosophila* genomes were accessed from NCBI (Coordinators, 2016; Drosophila 12 Genomes Consortium, 2007) —*D. simulans* (GCA_000259055.1), *D. sechellia* (GCA_000005215.1), *D. erecta* (GCA_000005135.1), *D. yakuba* (GCA_000005975.1), *D. ananassae* (GCA_000005115.1), *D. pseudoobscura* (GCA_000149495.1), *D. persimilis* (GCA_000005195.1), *D. willistoni* (GCA_000005925.1), *D. mojavensis* (GCA_000005175.1), *D. virilis* (GCA_000005245.1) and *D. grimshawi* (GCA_000005155.1)—while *D. melanogaster* (Release 6) was accessed from FlyBase (Goodman et al., 2019). Genomic scaffolds containing EcKLs were identified with TBLASTN searches (Altschul et al., 1990) using *D. melanogaster* EcKL protein sequences as queries and were annotated in Artemis (Carver et al., 2012). Gene model annotation was performed by reciprocally performing BLASTP and TBLASTN searches (Altschul et al., 1990) between putative gene models and *D. melanogaster* proteins with the highest sequence similarity. Annotated EcKL gene models in FlyBase in non-*D. melanogaster* species were used as starting points but were not assumed to be correct. EcKL domain completeness in translated gene models was checked with searches to the Pfam database (Punta et al., 2012). Gene models were considered pseudogenes if more than one inactivating mutation (frameshifts or premature stop codons) were found; gene models with only one inactivating mutation were considered null alleles, the stop codons of which were ignored for phylogenetic analyses or reverted to the most likely previous codon with a single nucleotide change. Gene models with missing exons due to incomplete genome assembly were considered “partial” genes and were assumed to be functional and not pseudogenes. Gene models were mapped to Muller elements (chromosome arms in *Drosophila*) using data from Schaeffer et al. (2008).

### 2.2. Phylogenetic analyses

Predicted protein sequences of EcKLs were aligned with MAFFT 7.402 (Katoh and Standley, 2013); proteins containing two domains were split into N- and C-terminal halves. MSAs were manually trimmed at the N- and C-termini to remove poorly-aligned regions using AliView (Larsson, 2014), and columns containing non-gap characters from only one sequence were also removed. Gene trees were generated with IQ-TREE v1.6.10 using ModelFinder to automatically pick the most appropriate model for the data (Kalyaanamoorthy et al., 2017; Nguyen et al., 2015), with the following command: *iqtree_1.6.10_comet -s infile.txt -bb 10000 -bnni -st AA -nm 50000 -msub nuclear -nt AUTO -pre output -m TESTNEW*. IQ-TREE was run five times on each alignment and the tree with the highest log-likelihood was used. MAFFT and IQ-TREE runs were conducted through the CIPRES Science Gateway (M. A. Miller et al., 2010).

A gene family clade was defined as a group of genes that all descended from a single gene inferred to exist in the most recent common ancestor of all 12 *Drosophila* species (MRCA_D_) based on parsimony, while a subclade was defined as a group of genes that all descended from a single gene inferred to be the product of a duplication event after the MRCA_D_, if that duplication event occurred in a common ancestor of at least two of the 12 *Drosophila* species. Clade and subclade nomenclature was developed to help refer to groups of orthologous genes, and individual genes have a clade ID in the format “DroC-S”, where C is the clade number and S is the subclade number. If a gene does not belong to a subclade, its subclade number is 0. This nomenclature is cladistic and based purely on inferred evolutionary relationships within the *Drosophila* genus, and clade IDs are not intended to be official gene names.

Ancestral clades were demarcated into four stability categories—stable (1:1 orthologs in every genome), blooming (at least one duplication, no complete losses), wilting (at least one complete loss, no duplications) and mixed (at least one duplication, at least one complete loss). For some downstream analyses, blooming, wilting and mixed clades were categorised as “unstable” clades. A gene loss event was defined as the loss (absence or pseudogenisation) of a gene lineage that existed in a species lineage after the MRCA_D_. A complete clade loss event was defined as the loss any of that clade’s genes in a particular species or lineage. A duplication event was defined as any inferred duplication of a gene that occurred after the MRCA_D_.

### 2.3. DGRP PheWAS

One hundred and forty six DGRP phenotypes were collected from 32 publications (Table S2) and aligned by DGRP line using the merge function in R (R Core Team, 2019). Abbott’s correction for control mortality was applied to methylmercury and caffeine phenotypes from Montgomery et al. (2014). DGRP genotypes, transcriptomes and annotations were recovered from the DGRP website (http://dgrp2.gnets.ncsu.edu/data.html). DGRP genotypes were first filtered for variants with a minor allele frequency > 0.05, and then for whether variants had been annotated to within—or within 1 kb of—EcKL genes or cytochrome P450 genes, resulting in 2,472 and 5,938 variants, respectively. Male and female transcriptomes were filtered for transcripts from EcKL or cytochrome P450 genes, including both copies of *Cyp12d1* found in the DGRP (*Cyp12d1-p* and *Cyp12d1-d*). PheWAS were performed by fitting a linear model in R between each variant or transcript and each DGRP phenotype. Genomic and transcriptomic associations were filtered using significance thresholds of *p* < 1×10^-5^ and *p* < 1×10^-3^ respectively.

### 2.4. Tissue-specific transcriptome data

Tissue-specific RNA-seq data in *D. melanogaster* were mined for the EcKL and P450 families from FlyAtlas2 (Leader et al., 2018). Tissue ‘enrichment’ was defined as (tissue FPKM + 1)/(whole body FPKM + 1) for each gene—this is roughly the ratio of tissue expression to whole body expression, but avoids undefined values and moves enrichment scores towards 1 for genes where the absolute difference between tissue and whole body FPKMs is small, effectively deweighting enrichment when expression measurement uncertainty is high and/or expression is less likely to be biologically relevant. Genes with enrichment values greater than or equal to 2 (expression ∼≥ 2-fold higher compared to whole body) were considered ‘enriched’ in a tissue, while genes with enrichment values less than 2 were considered ‘not enriched’.

### 2.5. Gene induction datasets

Seven xenobiotic differential gene expression datasets were mined for EcKL and P450 genes: phenobarbital in 3^rd^ instar larvae (W. W. Sun et al., 2006); phenobarbital in adult flies (King-Jones et al., 2006; Misra et al., 2011); piperonyl butoxide in adult flies (Willoughby et al., 2007); methamphetamine in adult flies (L. Sun et al., 2011); tunicamycin (8-hour timepoint) in adult flies (Chow et al., 2013); and fungal toxins (6-hour timepoint, wild-type vs. *ΔlaeA Aspergillus nidulans*) in 1^st^-instar larvae (Trienens et al., 2017). Genes were considered induced in a dataset if they were up-regulated ≥ 1.5-fold with a reported p-value < 0.05, except in the case of Chow et al. (2013), where their average up-regulation across all 20 lines needed to be ≥ 1.5-fold. Genes not meeting these criteria were considered not induced. Genes were considered induced (positively regulated) by CncC if they were up-regulated ≥ 1.5-fold (p < 0.05) upon ectopic expression of CncC in adult male flies, as reported by Misra et al. (2011), otherwise they were considered not induced.

### 2.6. Detoxification scores

*D. melanogaster* genes in the EcKL and P450 gene families were each given ‘detoxification scores’ from 0 to 4 based on four criteria (1 point was given for each criterion met): membership in a gene clade that is unstable in the *Drosophila* genus; induction in at least one xenobiotic induction dataset; induction by ectopic expression of CncC; and enrichment in at least one detoxification tissue (midgut, Malpighian tubules and/or fat body) at one or more life stages (3^rd^-instar larva, adult female and/or adult male). Data on the stability of P450 genes in *Drosophila* were taken from Good et al. (2014).

### 2.7. Review of published P450 functions

Published data on *D. melanogaster* P450 gene functions were collated from both FlyBase release FB2019_05 (Goodman et al., 2019) and direct searches for each gene name in the literature. Asserted gene functions were only recorded if they were supported with functional genetic or biochemical evidence (gene induction alone was not considered sufficient), and this evidence was rated as ‘weak’ (RNAi-only characterisation), ‘moderate’ (gene disruption and/or transgenic overexpression) or ‘strong’ (direct biochemical evidence, such as enzyme assays) depending on the experimental methods used. Unvalidated GWAS or TWAS associations with toxic stress phenotypes were not considered functional evidence for these purposes.

### 2.8. RNAi knockdown

*tubulin-GAL4*/TM3, *act-GFP*, *Ser^1^* females, which strongly express GAL4 in all tissues, were crossed to *UAS-dsRNA* males and the offspring were phenotypically scored for the presence or absence of the TM3, *act-GFP*, *Ser^1^* balancer chromosome. Significant deviation in genotypic ratios towards balancer-containing individuals was considered evidence for partial or complete developmental lethality associated with the inheritance of the *tubulin-GAL4* construct and the expression of the *UAS-dsRNA* construct. All fly crosses were conducted on yeast-cornmeal media at 25°C. dsRNA responder lines were obtained from the Vienna Drosophila Resource Center GD and KK libraries (www.vdrc.at; Table S5); KK lines were PCR genotyped for their hairpin landing site as per Green et al. (2014)—lines with annotated site insertions upstream of *tiptop* (which can produce dominant lethal effects) have been noted. The *tubulin-GAL4*/*actGFP*, *Ser^1^* line was a gift from Philip Batterham (The University of Melbourne). The 40D line was a gift from Kieran Harvey (Peter MacCallum Cancer Centre).

### 2.9. Statistical analyses

Interactions between xenobiotic induction, CncC induction and clade instability were modelled as a homogeneous association loglinear model with the *glm* function in R. Differential tissue enrichment between xenobiotic-induced/-uninduced, CncC-induced/-uninduced and stable/unstable genes was determined by comparing the median log_2_(enrichment) between groups of genes using bootstrap estimation with the *dabestr* package in R (Ho et al., 2019). Effect sizes with 95% confidence intervals that did not include 0 were considered significant. The diagnostic test for the sensitivity and specificity of the DS method was performed with the ‘diagnostic’ function in the *ThresholdROC* package (v2.7) in R. RNAi knockdown developmental lethality was assessed by comparing offspring genotypic ratios with *binom.test* in R, with a Bonferroni correction for multiple tests.

## 3. Results

### 3.1. Evolution of the EcKL gene family in the *Drosophila* genus

We explored the evolution of the EcKLs in the *Drosophila* genus by collating a dataset of gene models in the genomes of 12 species (Table S1). We manually annotated 564 EcKL gene models in the eleven non-*D. melanogaster* species in the *Drosophila* genus, to produce a dataset of 618 total EcKL gene models across the 12 species—of these, 605 are likely full gene models (of which 21 were considered likely pseudogenes), while 13 were partial gene models with clear missing sequence due to genome assembly incompleteness. The total number of putatively functional EcKLs (full and partial genes) in the genome of each species varies considerably; *D. willistoni* has the most with 61, while *D. mojavensis* has the least with 41 (Fig. 1). There are also differences between closely related species, the most striking one of which is the nine-gene difference between *D. simulans* (56 genes) and *D. sechellia* (47 genes). Two of these nine genes are missing from the assembly, while the remaining seven appear to have been pseudogenized in the *D. sechellia* lineage.

**Figure 1:**
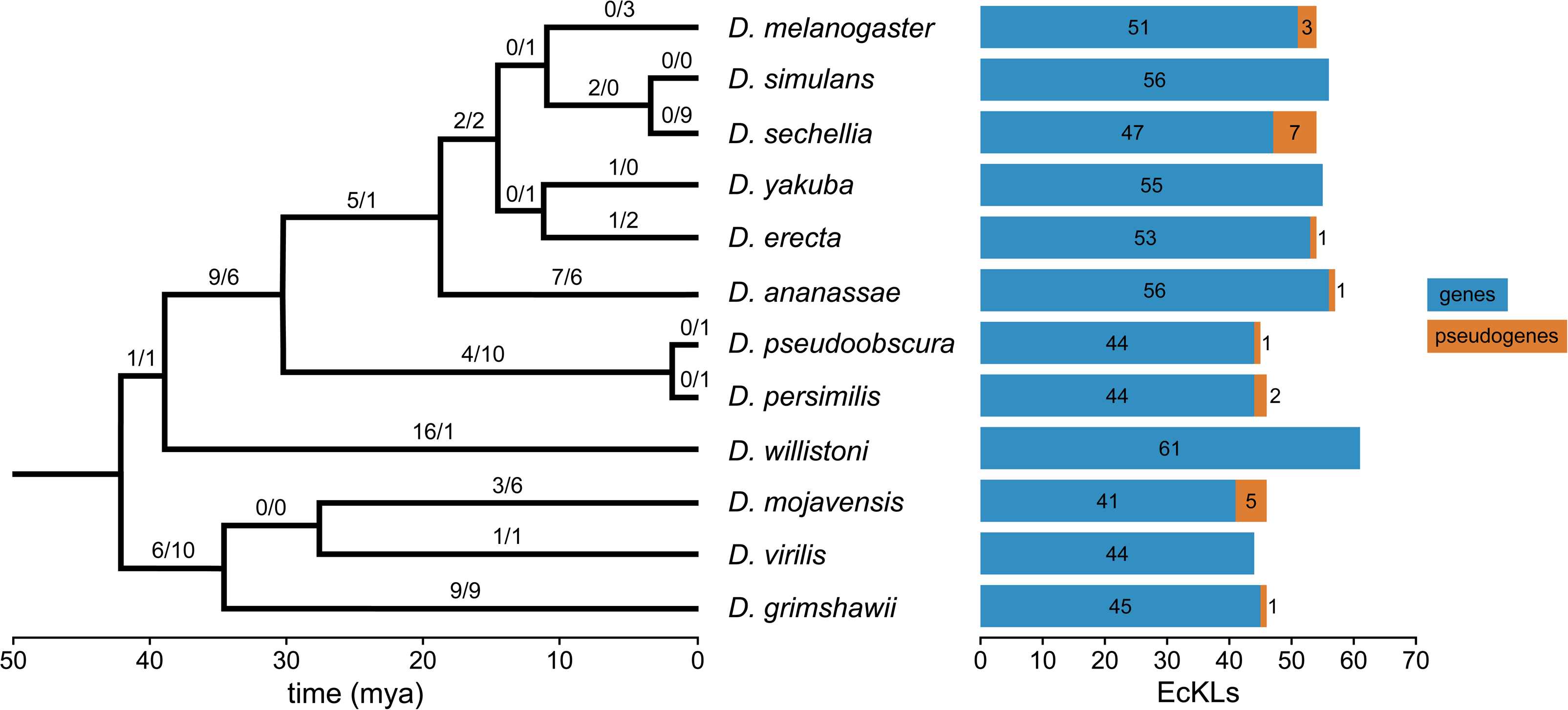
The number of EcKL genes (blue) and pseudogenes (orange) in the genomes of 12 *Drosophila* species. Numbers above the branches of the phylogenetic tree represent inferred gene gains and losses along that branch. Species tree from FlyBase (http://flybase.org/static/sequenced_species; Goodman et al., 2019).

By examining gene trees and using the known species phylogeny, we inferred the existence of 46 ancestral EcKL clades, which each represent a single EcKL that was present in the genome of the MRCA_D_. The vast majority of these clades have very strong support, however the reconstruction of the Dro26 clade is currently tentative and it may need to be split into smaller clades in the future. 18 clades (39%) are stable across the *Drosophila* genus, while 28 (61%) are unstable; of the unstable clades, 14 are blooming (at least one duplication after the MRCA_D_), 10 are wilting (at least one complete clade loss after the MRCA_D_) and 4 are mixed (at least one duplication and one complete clade loss after MRCA_D_; Fig. 2 and 3). The unstable clades are not all equally labile; while the mean number of inferred duplications and losses per unstable clade is 2.39 and 2.54 respectively, there are substantial outliers (Fig. 2). The Dro5 clade has experienced 20 duplications (and 10 losses) across the genus, and also has the highest number of genes in a single genome, with nine in *D. simulans*, while the Dro1 clade has the largest number of losses with 11

**Figure 2:**
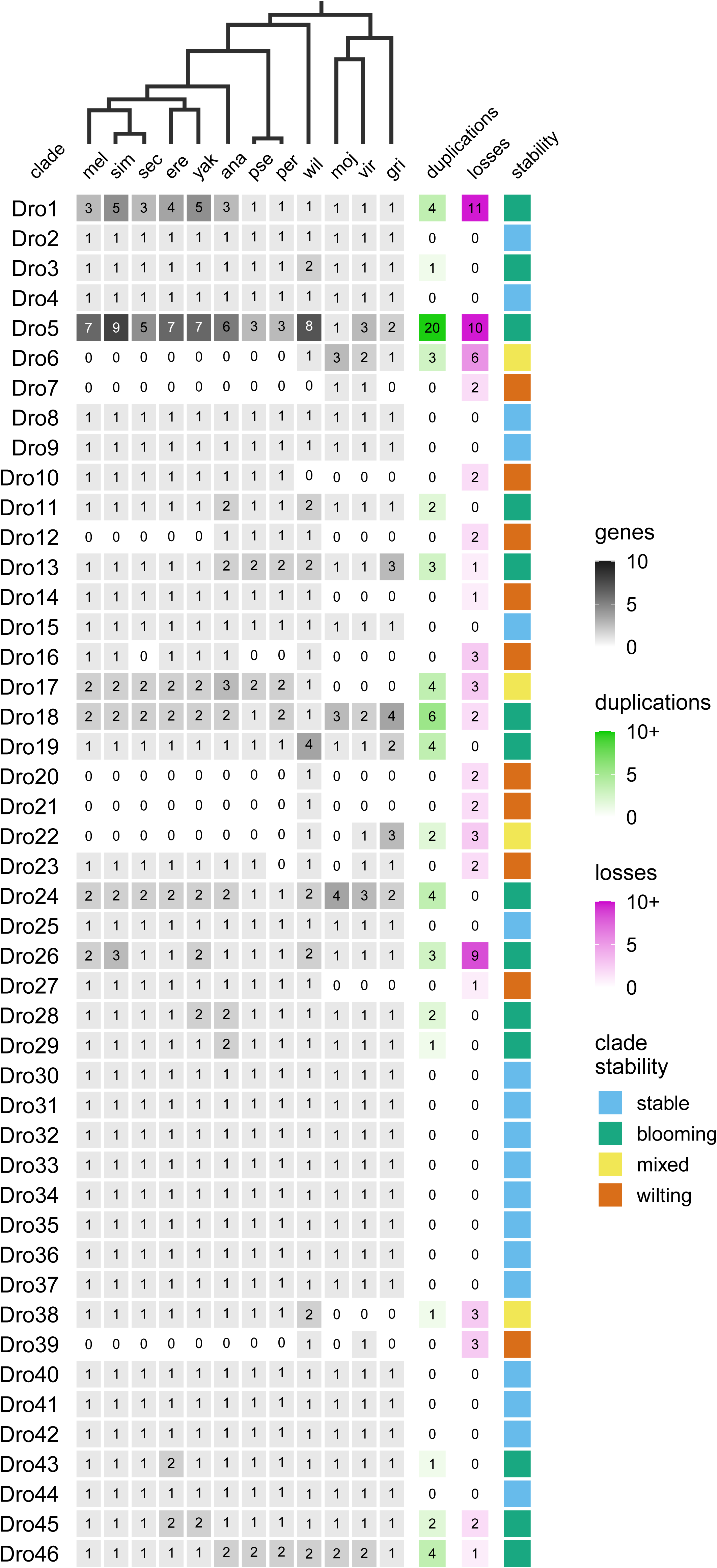
Number of EcKLs in each ancestral clade in each species of *Drosophila*, along with the inferred number of duplications and losses in—and the consequent stability of—each clade in the *Drosophila* genus.

**Figure 3:**
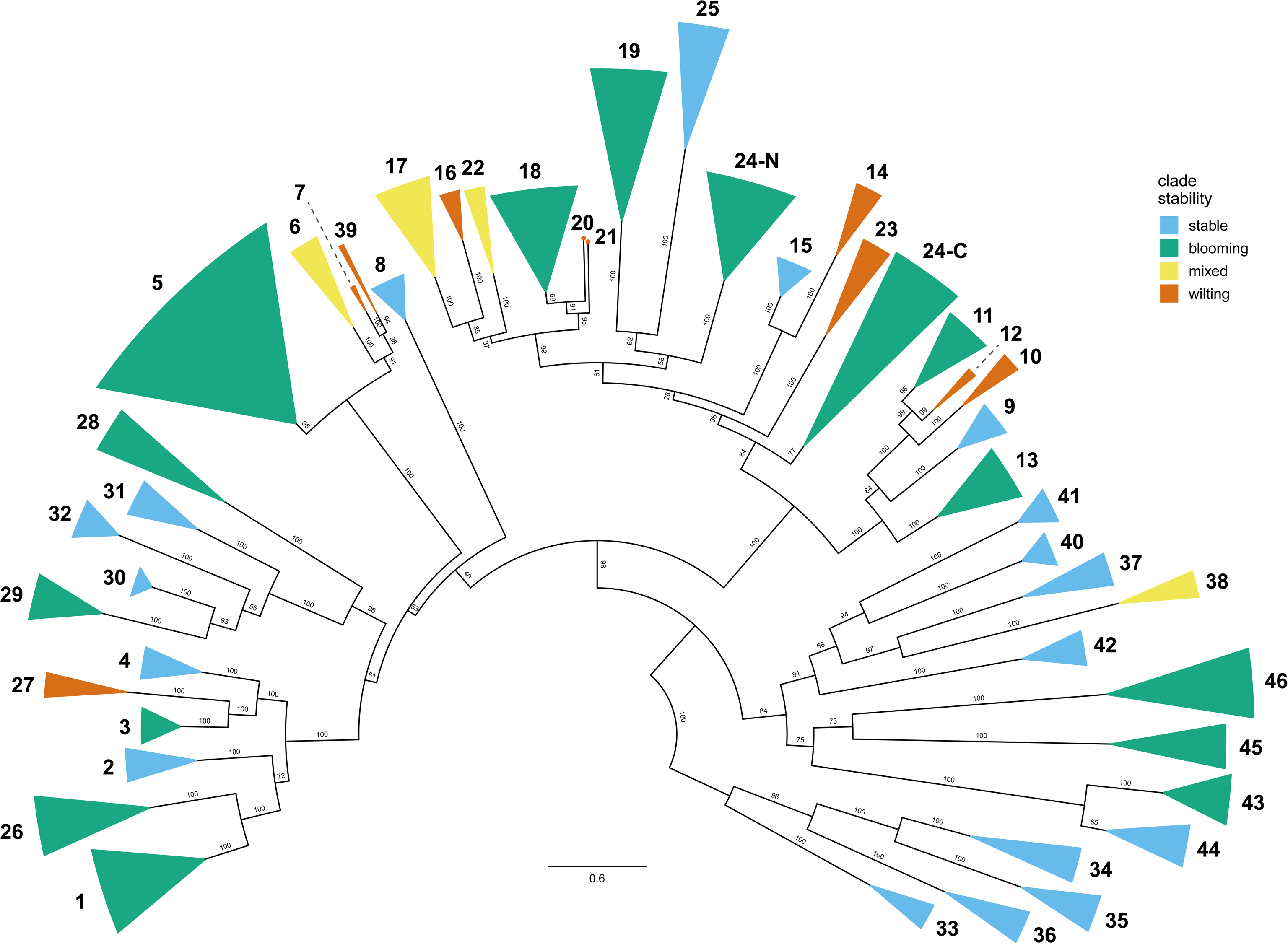
Gene tree of ancestral clades in the *Drosophila* genus, coloured by stability, and numbered by clade number; the tree was arbitrarily rooted on the branch leading to the Dro33, Dro34, Dro35 and Dro36 clades to facilitate its presentation, but the root has been omitted. Numbers on branches indicate bootstrap support values. The Dro24 clade is split into N- and C-terminal halves, 24-N and 24-C. The Dro20 and Dro21 clades comprise a single gene each, indicated by small circles.

We also found that EcKLs in *Drosophila* tend to cluster in uninterrupted tandem arrays—as is the case in other detoxification gene families (Feyereisen, 2011; Friedman, 2011; Robin et al., 1996)–the largest of which is on Muller element E (chromosome 3R in *D. melanogaster*) and contains 26 genes and two pseudogenes (encompassing all the genes in the Dro1 through Dro19 clades) in *D.* melanogaster, although this has been split into two clusters in the Drosophila subgenus, possibly by a chromosomal inversion or other rearrangement. This suggests tandem duplication is a significant cause of EcKL family expansion over time, and as many of these clusters contain genes from multiple clades, many of these duplications happened before the MRCA_D_. However, there are also instances of genes in the same clade found on different Muller elements, suggesting other processes, such as RNA-mediated gene duplication, transposition and translocation, may also contribute to gene family dynamics in the EcKLs.

### 3.2. EcKL and P450 genes are transcriptionally enriched in detoxification tissues in *D. melanogaster*

Detailed tissue-specific gene expression data in *D. melanogaster* published in FlyAtlas 2 (Leader et al., 2018) allowed us to explore the patterns of gene expression in both the EcKL and P450 families. FlyAtlas 2, at the time of writing, contains data from 18 tissues (head, eye, CNS, thoracicoabdominal ganglion, crop, midgut, hindgut, Malpighian tubules, fat body, salivary gland, trachea, ovary, virgin spermatheca, mated spermatheca, testis, accessory glands, carcass and rectal pad) and whole body samples, at between one and three life stages (3^rd^ instar larvae, adult males and adult females). Heatmaps of absolute expression (FPKM) for EcKLs and P450s can be found in Figs. S1 and S2, respectively. However, FPKM is a limited measure when comparing between genes, as some genes might require different levels of absolute expression to achieve similar functions (eg. the enzymes they encode may have different kinetic properties, or the transcribed or translated products may have their activities post-transcriptionally or post-translationally modified). As a more useful measure of tissue-specificity, we calculated the enrichment of each gene in each tissue using whole body expression at each life stage, where a greater enrichment value indicates more specific expression in that tissue, and an enrichment value of greater than or equal to 2 (or log_2_(enrichment) ≥ 1) indicates ‘enrichment’ (Figs. S3–4). As expected (Chung et al., 2009; J. Yang et al., 2007), many P450 genes are enriched in the midgut, Malpighian tubules and fat body at various life stages, which are widely considered detoxification tissues (Fig. 4A), but more EcKLs, proportionally, are enriched in these tissues, excepting the larval midgut: 10 (20%), 30 (59%) and 29 (57%) EcKLs are enriched in the midgut, and 30 (59%), 36 (71%) and 31 (61%) EcKLs are enriched in the Malpighian tubules, in larvae, adult females and adult males, respectively (Fig. 4B).

**Figure 4:**
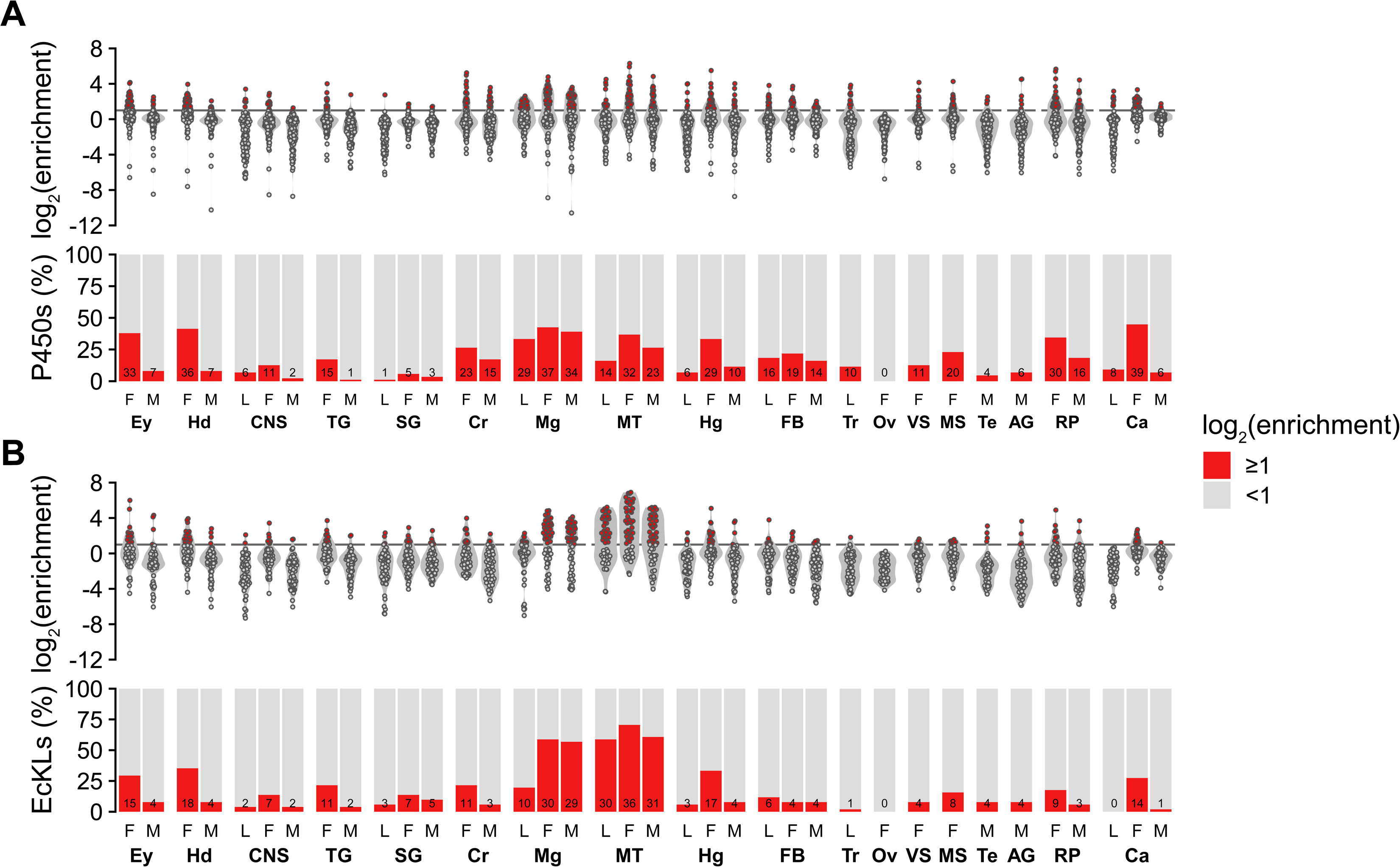
Tissue enrichment of P450 (A) and EcKL (B) genes (top) and the proportion of all genes in the family that are enriched (log_2_(enrichment) > 1; bottom) in *Drosophila melanogaster* across 18 tissues and three life stages. Horizontal lines on the top subfigure indicate a log_2_(enrichment) threshold of 1. Numbers in bars are the number of genes enriched in each tissue. L, 3^rd^ instar larva; M, adult male; F, adult female. Ey, eye; Hd, head; CNS, central nervous system; TG, thoracicoabdominal ganglion; SG, salivary gland; Cr, crop; Mg, midgut; MT, Malpighian tubules; Hg, hindgut; FB, fat body; Tr, trachea; Ov, ovary; VS, virgin spermatheca; MS, mated spermatheca; Te, testis; AG, accessory gland; RP, rectal pad; Ca, carcass.

### 3.3. Associations between xenobiotic induction, CncC induction, evolutionary stability and detoxification tissue enrichment

We used a collection of differential gene expression datasets to explore the transcriptional response of EcKL and P450 genes in *D. melanogaster* to the ingestion of xenobiotic compounds (phenobarbital, piperonyl butoxide, methamphetamine, tunicamycin and *Aspergillus nidulans* secondary metabolites) and to the ectopic expression of the xenobiotic-response factor CncC. We also used this data to test associations between the ‘detoxification properties’ of xenobiotic induction (defined as induction in at least one dataset), CncC induction and clade instability; we hypothesised that these two gene families will have significant groups of genes that have two or more of these properties and therefore that these properties will be associated gene family-wide. 34 (39%) P450s are induced in at least one xenobiotic dataset, compared with 23 (45%) EcKLs, while 20 (23%) P450s are induced by CncC, compared with 15 (29%) EcKLs (Fig. 5A,C). Family-wide, there are significant interactions between clade instability and xenobiotic induction (p = 0.005) and CncC induction and xenobiotic induction (p = 9×10^-4^) in the P450s (Fig. 5B); while in EcKLs, there is significant interaction only between clade instability and xenobiotic induction (p = 0.005; Fig. 5D).

**Figure 5:**
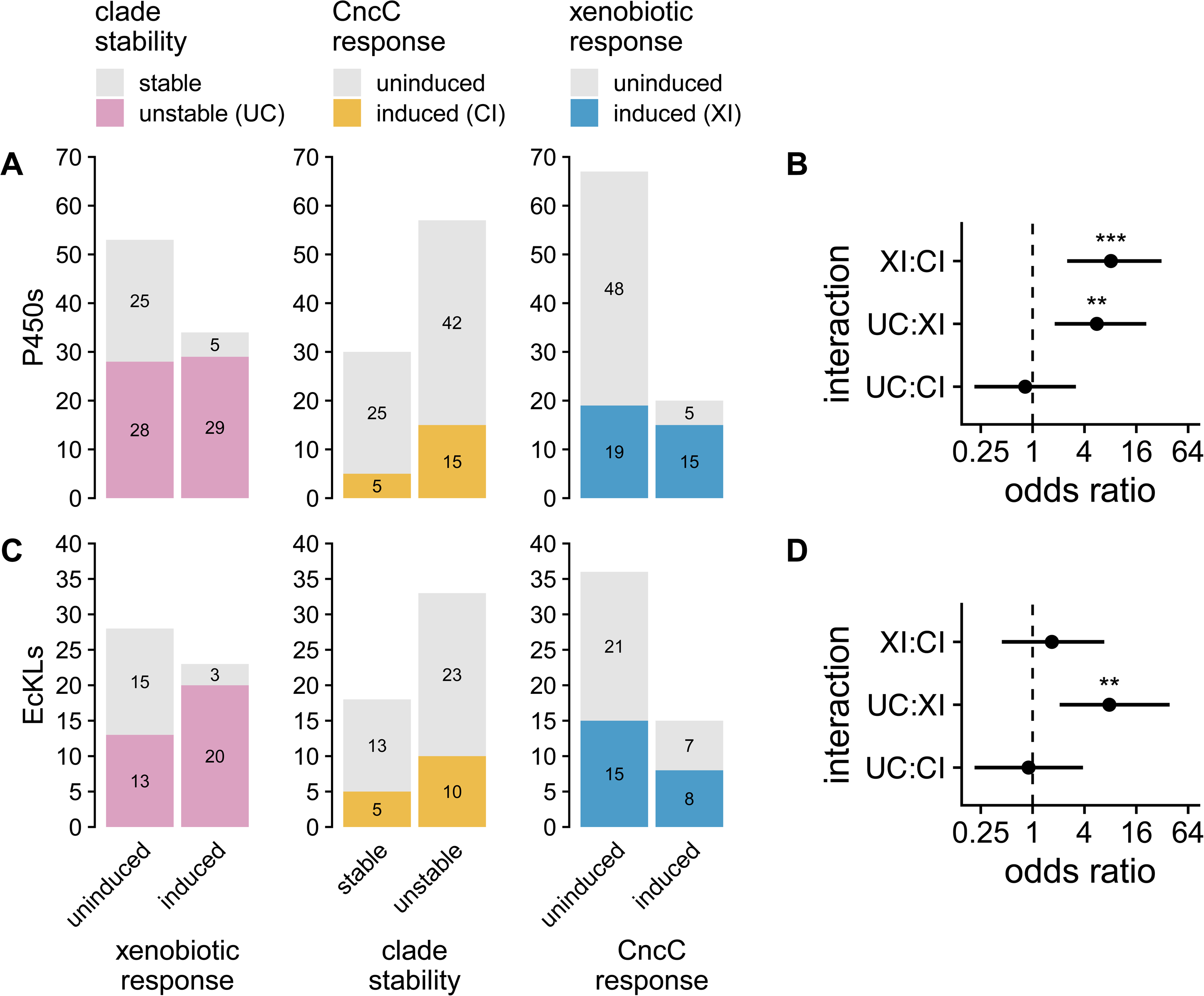
(A,C) The number of genes with sets of detoxification characteristics in *D. melanogaster*, for the P450s (A) and EcKLs (C). (B,D) Homogenous association loglinear models were fitted for the detoxification characteristics for both the P450s and the EcKLs. Odds ratios of interactions between pairs of characteristics for the P450s (B) and EcKLs (D) are shown; significant interaction is evidence of association between characteristics within the gene family. Errors bars are 95% confidence intervals. XI, xenobiotically induced; CI, CncC induced; UC, unstable clade. p-values: * < 0.05, ** < 0.01, *** < 0.001.

We then integrated these data with the tissue enrichment values calculated earlier to test if genes in each family with these detoxification properties have greater median enrichment in detoxification-related tissues than genes in the same family without these properties. P450s belonging to unstable clades have greater enrichment in the adult midguts and the adult female Malpighian tubules, while EcKLs belonging to unstable clades have greater enrichment in only the adult female midgut (Fig. S5). P450s induced by xenobiotics have greater enrichment in adult midguts and the adult female Malpighian tubules and fat body. In contrast, EcKLs induced by xenobiotics show greater enrichment in only the adult midguts and the adult female carcass (Fig. S6). P450s induced by CncC have greater enrichment in only the adult female Malpighian tubules, while EcKLs induced by CncC have greater enrichment in only the adult male Malpighian tubules (Fig. S7). Perhaps unsurprisingly, both EcKLs and P450s with detoxification properties tend to have reduced enrichment in a number of tissues not thought to play large roles in detoxification, including the larval CNS, salivary glands, hindgut and carcass, and the adult ovary and testes. However, we also noticed greater enrichment for EcKLs and P450s with detoxification properties in some tissues not typically associated with detoxification, such as the adult female eye, head, virgin and mated spermathecae and carcass, and the adult male carcass, particularly for the P450s (Figs. S5–7).

### 3.4. Genomic and transcriptomic variation in EcKL and P450 genes is associated with toxic stress phenotypes in *D. melanogaster*

A large amount of phenotypic, genomic and transcriptomic data is available for the Drosophila Genetic Reference Panel (DGRP), a collection of inbred lines of *D. melanogaster* that houses substantial naturally occurring genetic variation. Many of the phenotypes studied with the DGRP are related to toxic stress and can be used to detect candidate detoxification genes. We used a targeted phenome-wide association study (PheWAS) approach to detect associations between 146 phenotypes, including 55 toxic stress phenotypes (Table S2), and genomic and (adult sex-specific) transcriptomic variation in *D. melanogaster* P450 and EcKL genes, at a significance threshold of p < 10^-5^ for genomic variation, and p < 10^-3^ for transcriptomic variation. Summaries of PheWAS results can be found in Table S3A– F, while detailed outputs of the PheWAS analyses can be found in Table S4.

164 genomic variants in or near P450 genes were associated with 35 phenotypes, 26 of which related to toxic stress (Table S3A); most of these are linked together in haplotypes and have been previously reported in publications studying insecticide resistance (Battlay et al., 2018; 2016; Denecke et al., 2017; Duneau et al., 2018; L. Green et al., 2019; J. M. Schmidt et al., 2017)—however, some are novel. Four SNPs in or near *Cyp4d20* are associated with chlorantraniliprole survival; an intronic SNP in *Cyp4d8* is associated with DDT knockdown; a haplotype containing 10 linked SNPs in *Cyp4s3*, including a non-synonymous SNP (W260S), is associated with developmental caffeine survival; and a SNP producing a premature stop codon in *Cyp6a16*—which is annotated as a pseudogene in the *D. melanogaster* reference genome but rarely pseudogenized in natural populations (Good et al., 2014)—is associated with 3-hour malathion mortality in adult females. SNPs in or near five genes—*Cyp4ac1*, *Cyp4ac3*, *Cyp4c3*, *Cyp4e3* and *Cyp28d2*—were associated with ethanol tolerance in males and/or females. P450 transcript levels in males were associated with 36 phenotypes (25 toxic stress phenotypes) for 23 genes (Table S3B), while P450 transcript levels in females were associated with 37 phenotypes (29 toxic stress phenotypes) for 23 genes (Table S3C). Like the genomic variants, most of these have previously been reported. *Cyp4d8*’s transcript levels are associated with both boric acid and caffeine survival in adult females. *Cyp6a21* transcript levels in females are associated with developmental methylmercury survival.

70 genomic variants in or near EcKL genes were associated with 10 phenotypes, eight variants of which were associated with six toxic stress phenotypes (Table S3D); the bulk (56) of the total associations are part of the previously noted *CHKov1*-TE (Dro18-2) haplotype associated with sigma virus resistance (Magwire et al., 2011; 2012). A total of three SNPs near two paralogous but genetically unlinked EcKLs—*CG16898* (Dro26-1) and *CG33301* (Dro26-2)—are associated with developmental methylmercury survival, while a non-synonymous SNP (T4I) in *CG33301* is also associated with ethanol tolerance in adult females, and two downstream SNPs in *CG33301* are associated with larval activity during exposure to the neonicotinoid imidacloprid. EcKL transcript levels in males were associated with 10 phenotypes (five toxic stress phenotypes) for nine genes (Table S3E), while transcript levels in females were associated with 10 phenotypes (seven toxic stress phenotypes) for five genes (Table S3F). Many of these were associations between *CG6908* (Dro23-0) and chlorantraniliprole and malathion survival, which is an unvalidated candidate gene for resistance to these two insecticides. *CG10560* (Dro17-3) expression is also associated with chlorantraniliprole survival in males and females, while *CG11878* (Dro1-5) expression in females is associated with ethanol tolerance in males.

### 3.5. The detoxification score method is concordant with known functions of *D. melanogaster* P450s

As detoxification genes have generally accepted evolutionary and transcriptomic characteristics, we developed ‘detoxification score’ (DS) that scores genes (0–4) based on the criteria of evolutionary instability, xenobiotic induction, CncC induction and detoxification tissue enrichment. Genes with scores of 3 or 4 were considered detoxification candidates. We tested the accuracy of the DS method by applying it to the P450 gene family in *D. melanogaster* (Fig. 5) and comparing the scores with published functions of 31 P450 genes (Table 2).

**Table 2:**
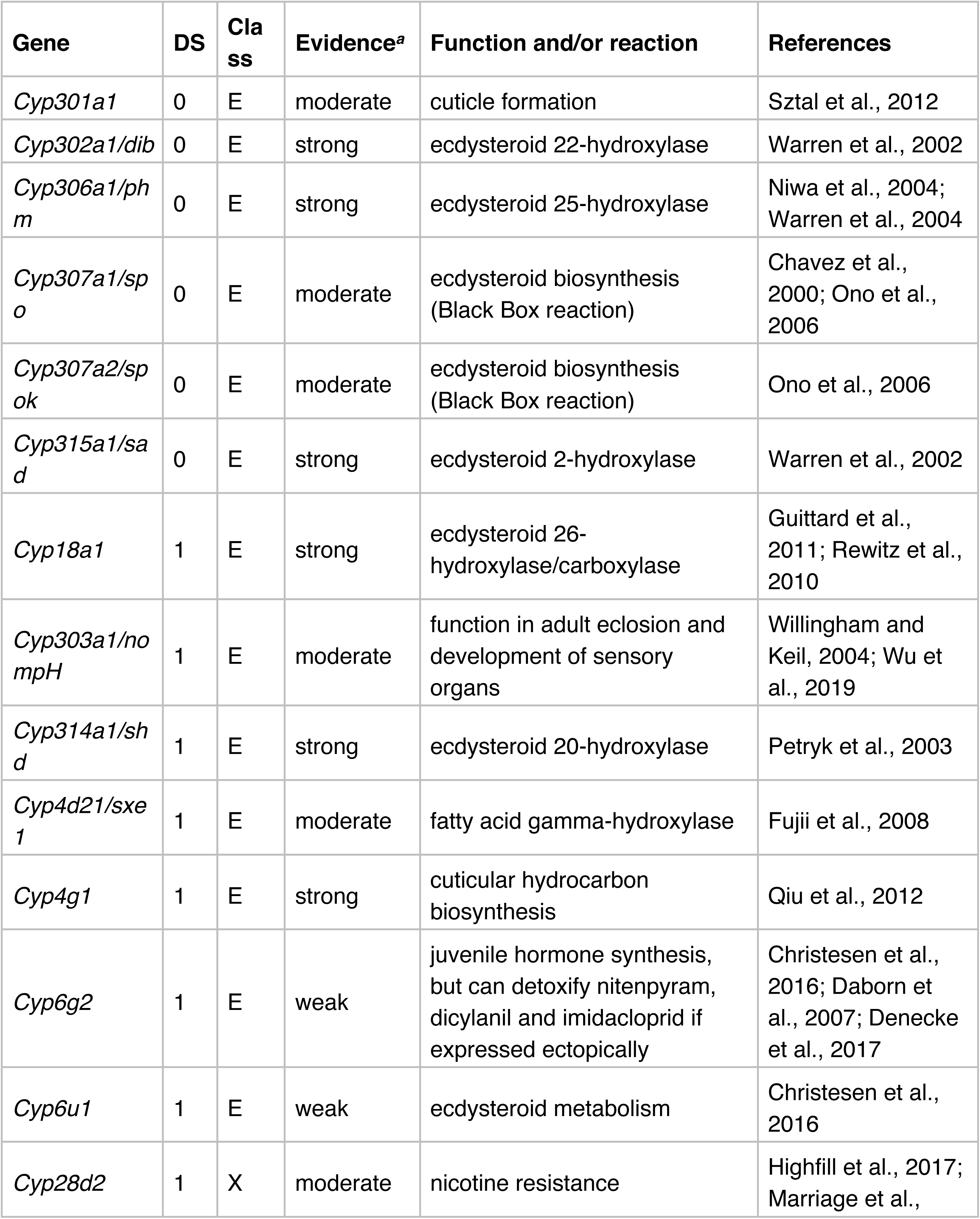
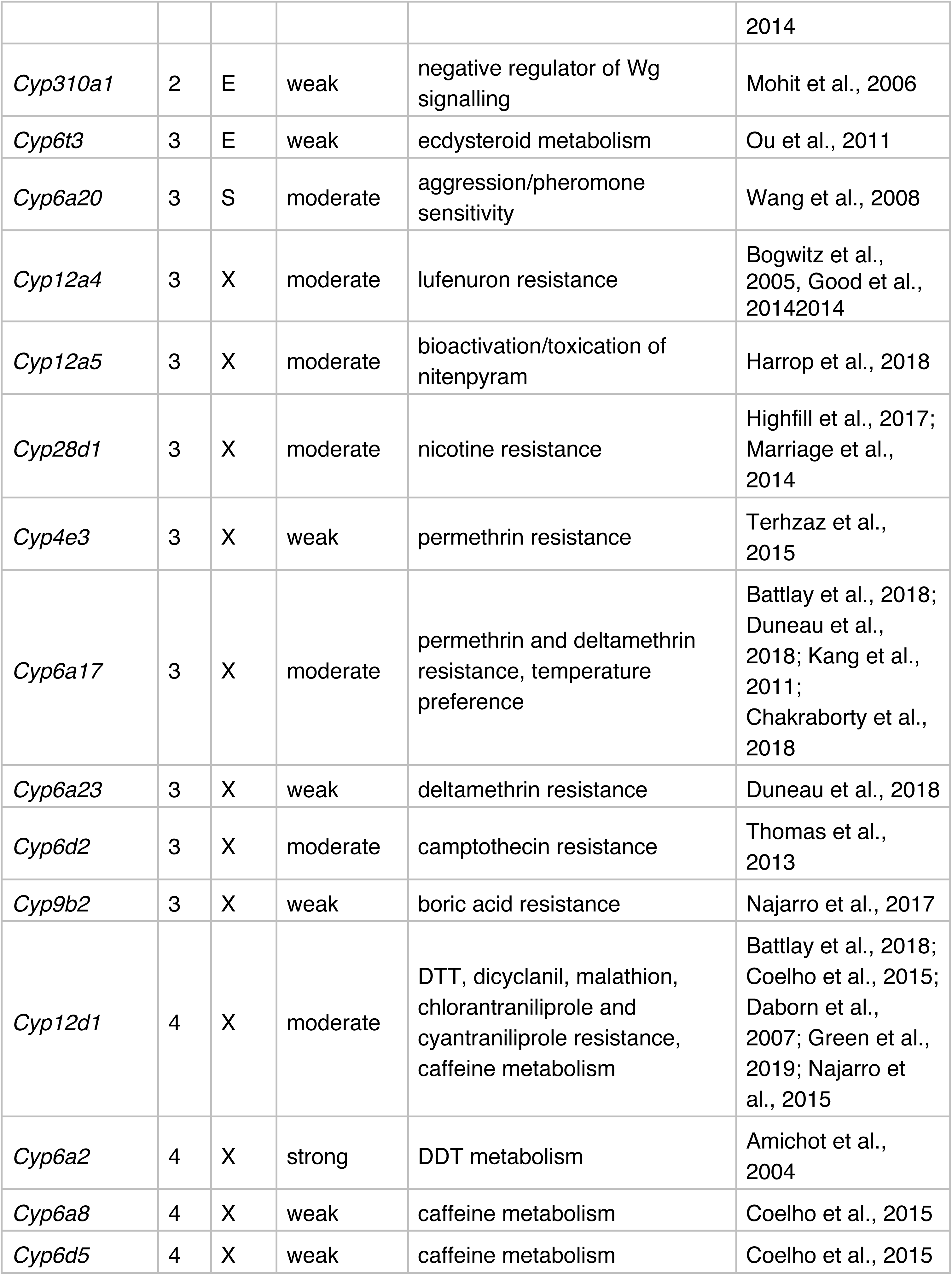
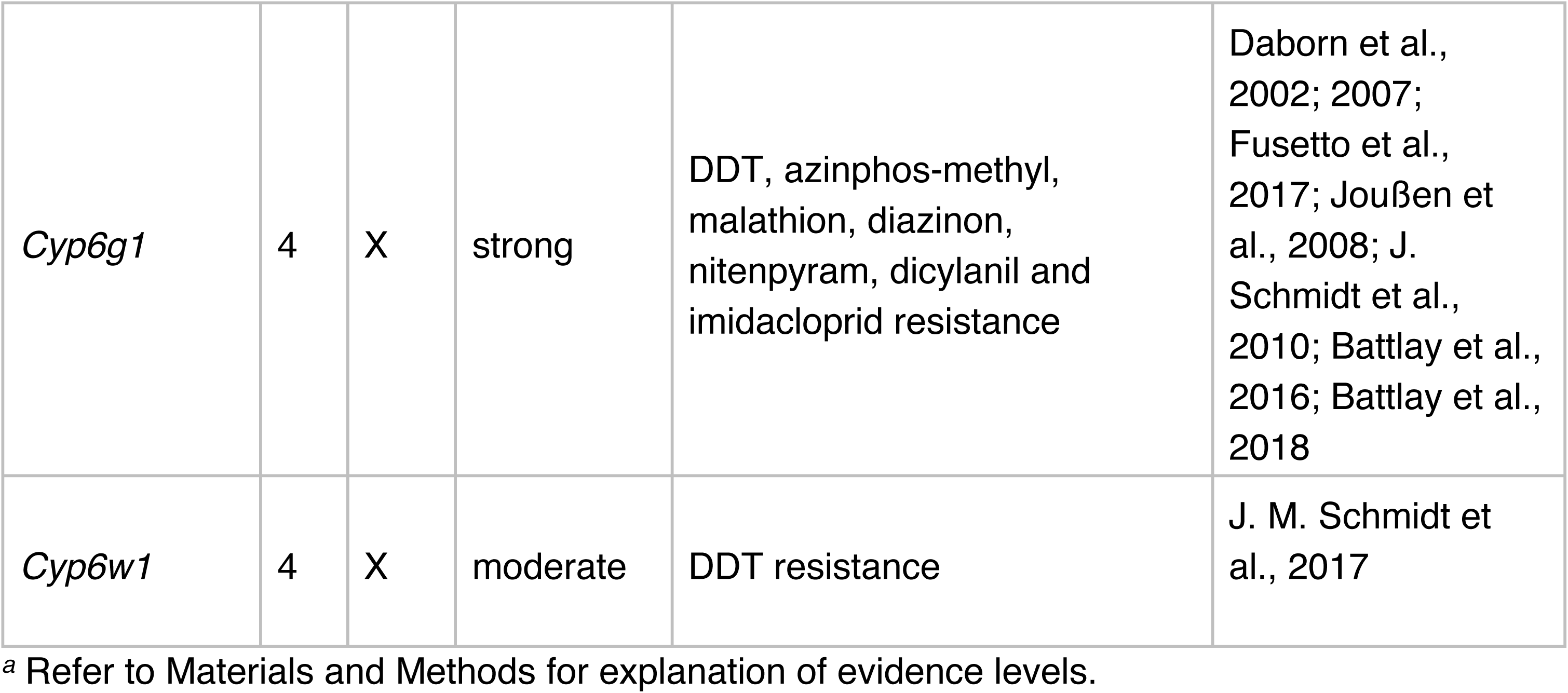
Comparison of detoxification score (DS) values with known P450 functions in *Drosophila melanogaster*.

Fifteen P450s have X-class (detoxification) functions described in the literature, of which only one, *Cyp28d2* (DS = 1), had a score less than 3. *Cyp28d2* has been implicated in nicotine resistance, along with its paralog *Cyp28d1* (DS = 3; Highfill et al., 2017; Marriage et al., 2014). The evidence for both genes’ roles in nicotine detoxification comes from quantitative complementation with loss-of-function alleles, as well as ubiquitous RNAi knockdown (Highfill et al., 2017), but they have not been characterised further. *Cyp12a5* (DS = 3) is another interesting case, given that it has been implicated in the bioactivation—not detoxification—of the neonicotinoid insecticide nitenpyram (Harrop et al., 2018). However, given that it putatively metabolises a xenobiotic compound, even though that metabolism increases its toxicity, it seems likely that it detoxifies other similar compounds. *Cyp6a17* (DS = 3) has characterised roles in pyrethroid insecticide resistance (Battlay et al., 2018; Duneau et al., 2018), but also has a connection to temperature preference behaviour (Chakraborty:2018fh; Kang et al., 2011). This is not the only behavioural phenotype linked to a P450: *Cyp6a20* (DS = 3), has a putatively S-class (secondary metabolic) function due to its links to male aggression (Dierick and Greenspan, 2006; Robin et al., 2007; L. Wang et al., 2008).

Fifteen P450s have E-class (essential physiological or developmental) functions described in the literature, of which only one, *Cyp6t3* (DS = 3), had a score greater than 2. *Cyp6t3* has a proposed role in ecdysteroid biosynthesis based on RNAi evidence (Ou et al., 2011), but the gene has undergone a complete loss event in the *Drosophila* genus (Good et al., 2014), which is inconsistent with a conserved function in hormone metabolism. Nevertheless, *Cyp6t3* is not enriched in any of the three detoxification tissues and so may not be a detoxification gene. Additionally, *Cyp6g2* (DS = 1) is an interesting case, as its proposed role in juvenile hormone synthesis (Christesen et al., 2016) and corpora allata-specific expression (Chung et al., 2009) are at odds with its ability to detoxify multiple insecticides when ectopically expressed in detoxification tissues (Daborn et al., 2007; Denecke et al., 2017).

Based on these data, the accuracy of the DS method was evaluated with a diagnostic test. The sensitivity (true positive rate) was 93.3% (95% CI: 68.1–99.83), while the specificity (true negative rate) was 87.5% (95% CI: 61.65–98.45). Considering the small sample sizes available, this suggests the DS method is reasonably accurate at predicting known detoxification functions of P450s and has the potential to identify candidate detoxification genes for follow-up analyses. Indeed, 16 P450s that had scores of 3 or 4 have not yet been characterised—we predict that between 11–16 of these genes encode detoxification enzymes (Table 3).

**Table 3:** *Drosophila melanogaster* P450 genes with detoxification scores (DS) greater than 3 and with currently unknown functions.

| Detoxification score | Genes |
| --- | --- |
| 3 | *Cyp4ac3, Cyp4d1, Cyp4d14, Cyp4p2, Cyp4s3, Cyp6a13, Cyp6a14, Cyp6d4, Cyp6t1, Cyp12c1, Cyp28a5* |
| 4 | *Cyp4d2, Cyp4e2, Cyp4p1, Cyp6a21* |

### 3.6. DS method suggests many *D. melanogaster* EcKLs are candidate detoxification genes

Applying the DS method to the EcKL family in *D. melanogaster* (Fig. 7), 24 genes (47%) have a DS ≥ 3 (Fig. 8B), compared with 32 genes (36.8%) in the P450 family (Fig. 8A). Based on the aforementioned sensitivity of the DS method that used the P450 family as a ‘truth set,’ we estimate between 16–24 EcKLs are involved in detoxification. Seven EcKLs have a DS of 4: *CG31288* (Dro11-0), *CG10550* (Dro13-1), *CG10553* (Dro17-1), *CG10560* (Dro17-3), *CG10562* (Dro18-1), *CG6908* (Dro23-0) and *CG9498* (Dro29-0). Interestingly, *CG10562* is a direct paralog of *CHKov1* (Dro18-2, DS = 2), which has been implicated in resistance to sigma virus infection (Magwire et al., 2012; 2011). Five and two genes in the highly unstable Dro5 and Dro26 clades, respectively, were also detoxification candidates (DS = 3).

**Figure 7:**
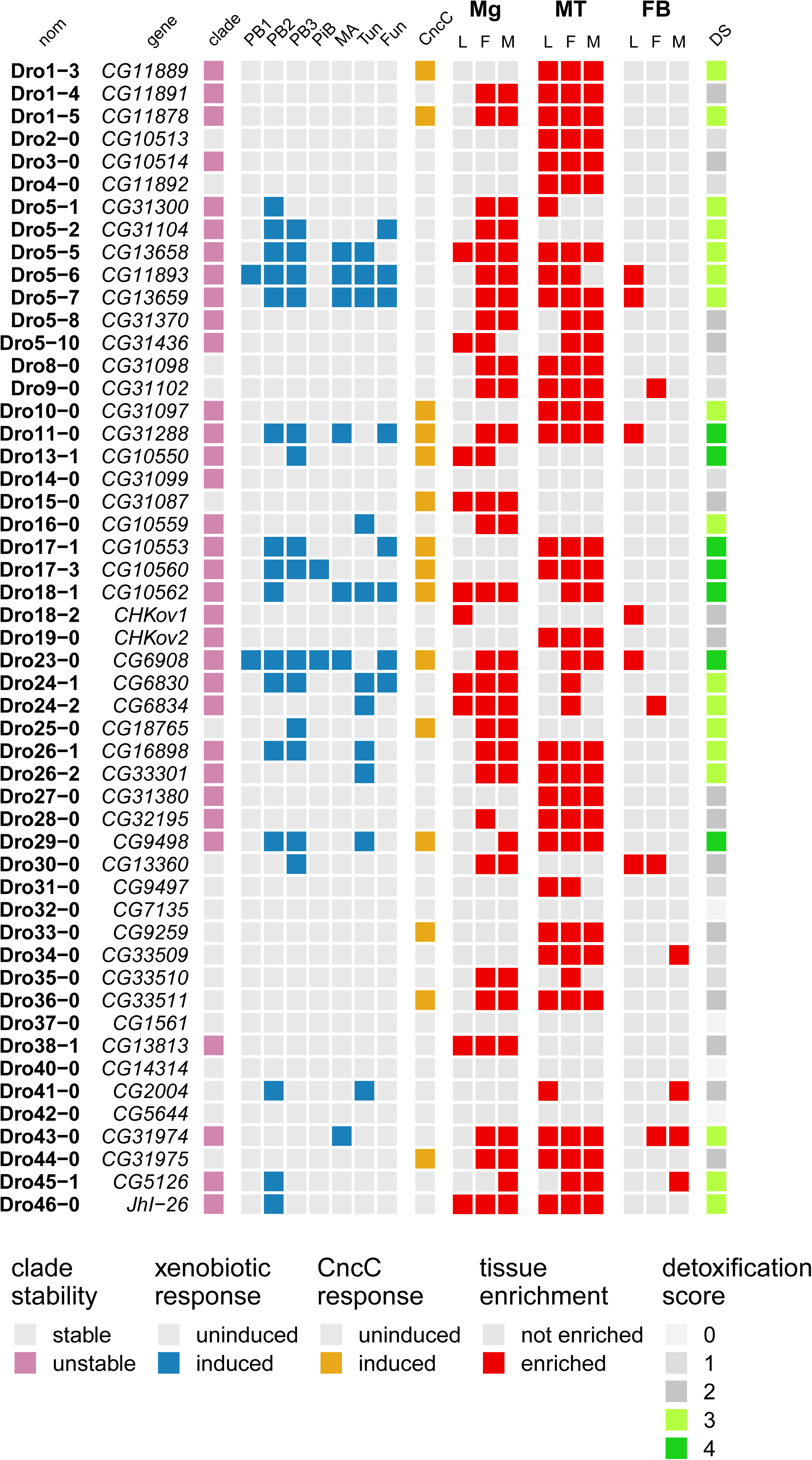
Application of the detoxification score (DS) to EcKL genes in *Drosophila melanogaster*, wherein genes receive one point for meeting each detoxification criterion, indicated by non-grey coloured squares. Genes are identified by both their *Drosophila* ancestral clade nomenclature, and their name or annotation symbol, as appropriate. PB1, phenobarbital dataset 1 (W. W. Sun et al., 2006); PB2, phenobarbital dataset 2 (King-Jones et al., 2006); PB3, phenobarbital dataset 3 (Misra et al., 2011); PiB, piperonyl butoxide (Willoughby et al., 2007); MA, methamphetamine (L. Sun et al., 2011); Tun, tunicamycin (Chow et al., 2013); Fun, *Aspergillus nidulans* toxins (Trienens et al., 2017); Mg, midgut; MT, Malpighian tubules; FB, fat body; L, 3^rd^ instar larvae; F, adult females; M, adult males; DS, detoxification score.

**Figure 8:**
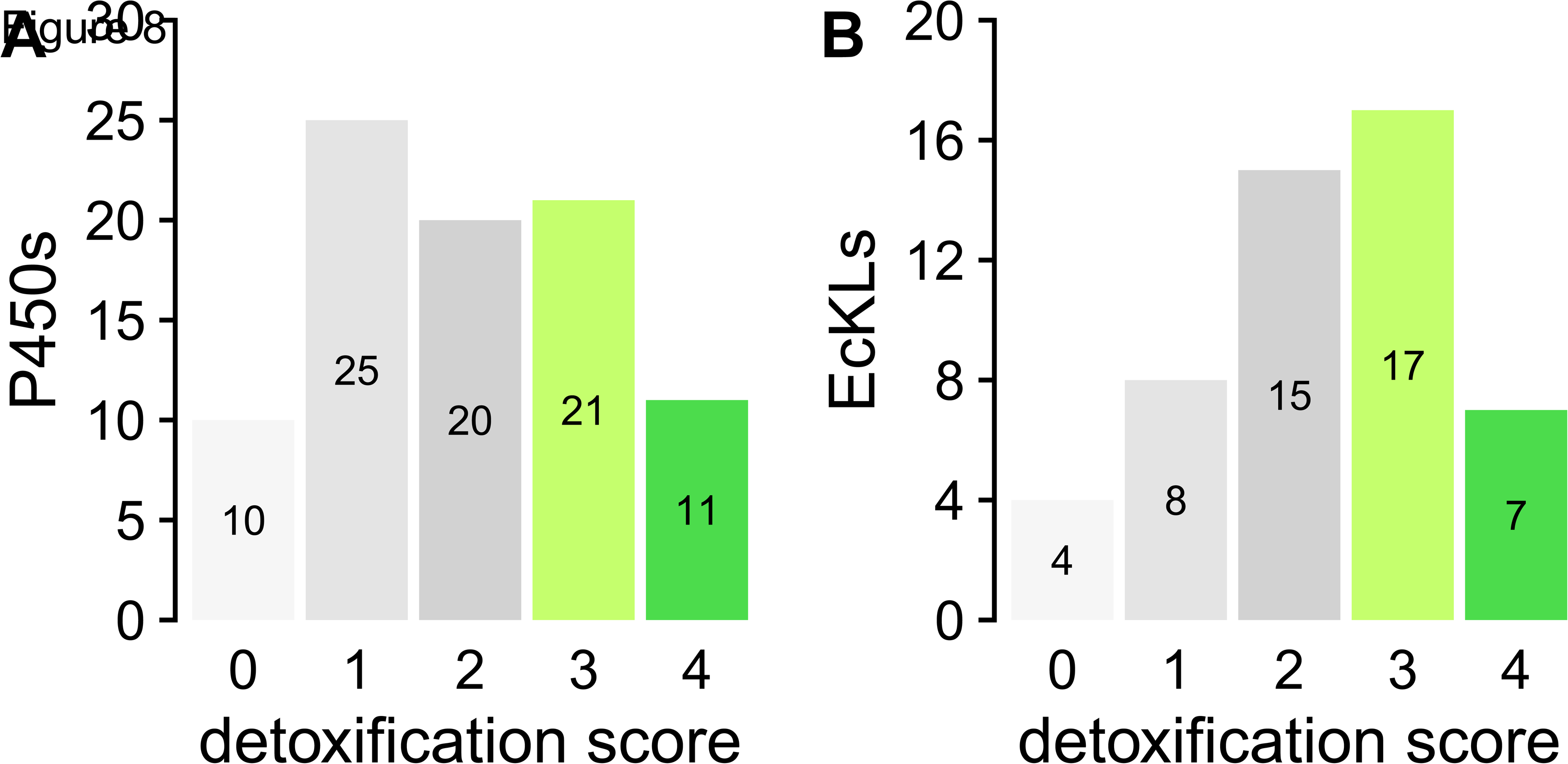
The number of genes in the P450 (A) and EcKL (B) gene families in *Drosophila melanogaster* with a given detoxification score. Scores of 3 and 4 indicate candidate detoxification genes.

### 3.7. Ubiquitous RNAi knockdown of EcKLs in *D. melanogaster*

As a preliminary attempt to find EcKLs essential for development, we crossed UAS-dsRNA responder males for 41 EcKLs (and three control lines) to females containing a *tub*-GAL4 driver over a balancer chromosome, and phenotypically scored the eclosing adult offspring (Fig. S8). 14 of the 41 EcKL knockdown crosses produced significantly fewer putative knockdown individuals than expected, indicating some degree of developmental lethality. However, six of these crosses involved lines from the KK library containing annotated hairpin insertions, which can produce developmental defects due to ectopic expression of the gene *tiptop* and activation of the Hippo pathway (E. W. Green et al., 2014; Vissers et al., 2016). A control cross with the 40D responder line, which contains a UAS-only insertion at the annotated site, produced very few driver-containing offspring, strongly suggesting that the developmental lethality observed with the six annotated KK lines is due to *tiptop* misexpression and not the knockdown of the EcKL targeted by the dsRNA. Overall, only eight EcKLs resulted in significant developmental lethality that was attributable to their reduced expression: *CG31102* (Dro9-0), *CG31099* (Dro14-0), *CG10562* (Dro18-1), *CHKov1* (Dro18-2), *CG9497* (Dro31-0), *CG13813* (Dro38-1), *CG2004* (Dro41-0) and *CG31975* (Dro44-0).

## 4. Discussion

### 4.1. EcKL evolution in *Drosophila*

The evolutionary pattern seen in the EcKL gene family this study matches that seen in canonical detoxification gene families in *Drosophila*, such as the P450s, glutathione S-transferases (GSTs) and carboxylcholinesterases (CCEs): a mix of stable clades that have 1:1 orthologs in all species, and unstable clades with high rates of gene gain, gene loss or both (Good et al., 2014; Low et al., 2007; Robin et al., 1996; 2009). We have found that a few EcKLs belonging to stable clades in *Drosophila* are also highly conserved across insects as a whole (Scanlan and Robin, unpublished); these are good candidates for genes with essential E-class functions. It is important to note, however, that not all genes with E-class functions are necessarily stable over evolutionary time: *Cyp4g1* and *Cyp4g15* orthologs in insects encode enzymes responsible for alkane and alkene biosynthesis, yet this physiologically essential function is maintained despite seemingly random duplications and losses in many lineages (Feyereisen, 2020); similarly, orthologs of the Halloween genes *spook* (*Cyp307a1*) and *spookier* (*Cyp307a2*) encode genes essential for the biosynthesis of ecdysteroids, yet also have experienced elevated rates of duplication and loss in insects and other arthropods (Perry et al., 2019; Rewitz et al., 2007; Sezutsu et al., 2013; Sztal et al., 2007).

Of interest in this study is the nine-gene difference in EcKL number between *D. simulans* and *D. sechellia*, which is due to gene losses in the latter species in only four clades: Dro5 (four losses), Dro1 (two losses), Dro26 (two losses) and Dro16 (one loss). In *D. melanogaster*, many of the genes in these clades are detoxification candidates, suggesting genes in these clades in other species may also function in detoxification. *D. sechellia* also has fewer P450 genes compared to *D. simulans*—74 versus 88 (Good et al., 2014)—and gene losses have also been observed in the glutathione S-transferase and odorant receptor gene families in this species (Low et al., 2007; McBride et al., 2007). Two competing hypotheses have been proposed to explain this concerted gene loss: the first is that moving from a generalist niche, as seen with *D. simulans*, to a specialist niche feeding on morinda fruit (Dworkin and Jones, 2008) has led to relaxed selection on genes involved in detoxification and olfaction, producing an elevated rate of pseudogenisation in the *D. sechellia* lineage; the second is that possible severe population reduction after the divergence of *D. sechellia* and *D. simulans* resulted in elevated fixation of slightly deleterious null alleles (Good et al., 2014). If the first is true, this may be further evidence for the hypothesised detoxification functions of genes in the Dro1, Dro5, Dro16 and Dro26 clades.

A limitation of this study of EcKL evolution is that the genome assemblies used, apart from that of *D. melanogaster*, are of relatively poor quality compared to more recently published assemblies (D. E. Miller et al., 2018). The BUSCO scores (which indicate the percentage of conserved single-copy orthologs in a genome; a measure of assembly completeness) of the assemblies used here range from 96.6% to 98.6% (D. E. Miller et al., 2018), suggesting some orthologs annotated as “lost” in this study may simply be missing from the assembly but not the genome. Future work on EcKL evolution should endeavour to use the highest quality genome assemblies available to minimise this issue.

### 4.2. Detoxification gene properties and tissue-specific enrichment

Detoxification genes are thought to have various properties, including evolutionary instability, transcriptional induction by toxins and toxin-response pathways, and enrichment in detoxification-related tissues (Kawashima and Satta, 2014; Misra et al., 2011; J. H. Thomas, 2007; Willoughby et al., 2006; J. Yang et al., 2007). If this is true, these properties should associate together in detoxification families, but to the best of our knowledge this has yet to be tested in any detail. The results in this study generally support this hypothesis, with some detoxification characteristics associating in the P450s, a canonical detoxification gene family.

The tissue enrichment data analysed here show EcKLs are strongly enriched in the Malpighian tubules, with comparatively little enrichment (proportional or otherwise) in the larval midgut (Fig. 4B). While both tissues are thought to be involved in detoxification, it is unclear how the detoxification processes differ between the midgut and the Malpighian tubules. A reasonable assumption is that the midgut is involved in immediate “first-pass” xenobiotic metabolism—similar to the mammalian small intestine and liver (Kaminsky and Zhang, 2003; Pond and Tozer, 1984)—while the Malpighian tubules are involved in reactions that promote the excretion of xenobiotic compounds, such as conjugations, and detoxify compounds that are already circulating in the hemolymph—similar to the mammalian liver and kidney (Bajaj et al., 2018; Lock and Reed, 1998). Detoxicative kinases, while conceptually catalysing a phase II conjugation reaction, may also act on toxins in the midgut directly if they contain hydroxyl groups, or on the immediate products of P450-mediated hydroxylation. Interplay between detoxification reactions within and between tissues is a topic that deserves greater study in insects.

It is important to note that the tissue expression data from FlyAtlas 2 (Leader et al., 2018) considered in this paper consist of basal levels of expression, and given that xenobiotic compounds may induce expression of detoxification genes—particularly in detoxification tissues (Willoughby et al., 2006)—the tissue-specific enrichment of detoxification genes during xenobiotic exposure is likely to be different to what is observed here. Comparatively low levels of basal enrichment of EcKLs in the larval midgut might be offset by extensive midgut-specific induction of these genes upon exposure to relevant toxins. In contrast, we hypothesise that the high levels of basal enrichment of EcKLs in adult midguts might be due to relatively high constitutive expression of detoxification genes in these tissues, which may be adaptive for the intermittent feeding behaviour of adults (Xu et al., 2008). The FlyAtlas 2 data used here are also derived from a single fly line, Canton S, which might show transcriptomic differences with other lines; however, our DS method can reliably detect detoxification genes identified in other genetic backgrounds (Table 2), suggesting the method is robust across genotypes.

A drawback of analysing tissue enrichment at a family-wide level is that it elides differences between genes—some detoxification genes have highly tissue-specific expression patterns (ie. expressed in the midgut but not the Malpighian tubules, or vice versa; (J. Yang et al., 2007). Some non-detoxification genes may also have high enrichment in these “detoxification” tissues—this may partially explain the large confidence intervals seen for differential Malpighian tubule enrichment in the EcKLs (Figs. S6–8), as some genes with extremely high enrichment in this tissue (eg. *CG10513*, *CG10514*, *CG11892* and *CG9259*) do not display other detoxification characteristics and consequently are poor detoxification candidates (Fig. 6).

**Figure 6:**
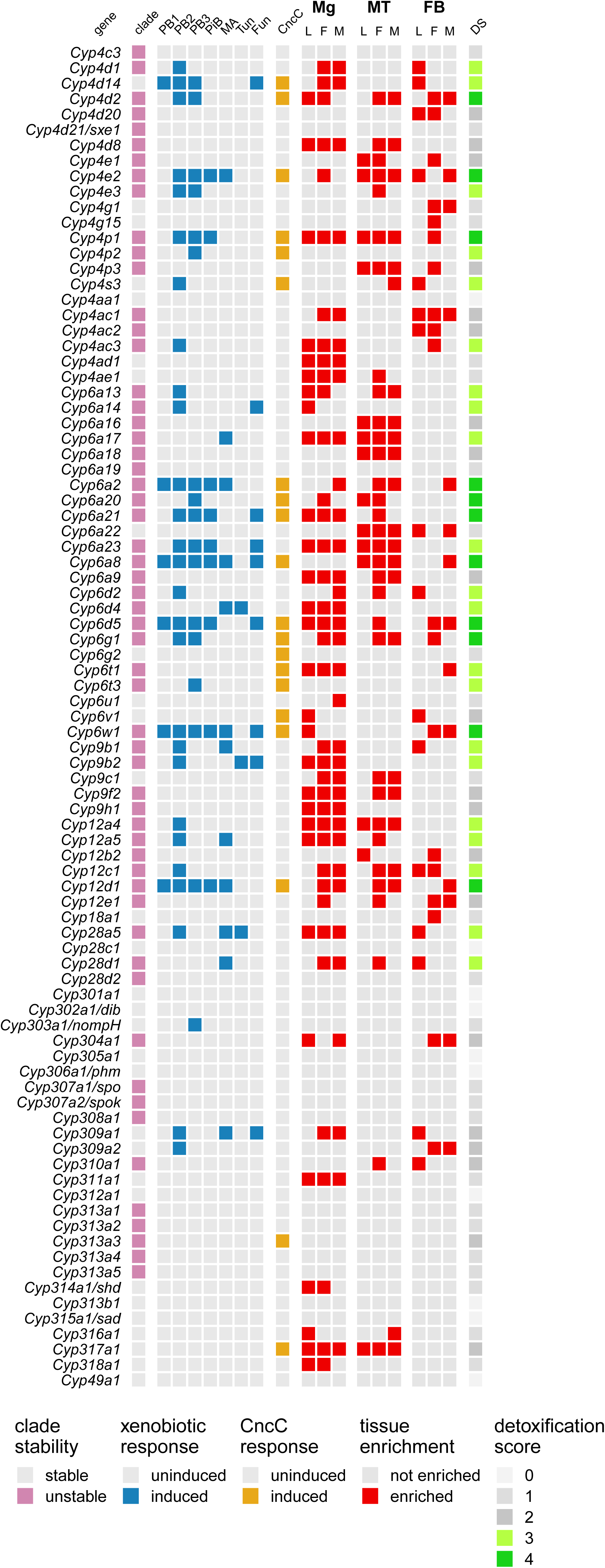
Application of the detoxification score (DS) to P450 genes in *Drosophila melanogaster*, wherein genes receive one point for meeting each detoxification criterion, indicated by non-grey coloured squares. Genes with names other than P450 nomenclature designation have both indicated, separated by a slash. PB1, phenobarbital dataset 1 (W. W. Sun et al., 2006); PB2, phenobarbital dataset 2 (King-Jones et al., 2006); PB3, phenobarbital dataset 3 (Misra et al., 2011); PiB, piperonyl butoxide (Willoughby et al., 2007); MA, methamphetamine (L. Sun et al., 2011); Tun, tunicamycin (Chow et al., 2013); Fun, *Aspergillus nidulans* toxins (Trienens et al., 2017); Mg, midgut; MT, Malpighian tubules; FB, fat body; L, 3^rd^ instar larvae; F, adult females; M, adult males; DS, detoxification score.

### 4.3. Suitability of DS method for identifying candidate detoxification genes in insects

In this study, we have proposed a “detoxification score” (DS) for the identification of candidate detoxification genes from a combination of evolutionary and transcriptomic data available in *D. melanogaster*. We chose an integrative approach because each individual detoxification property (evolutionary instability, xenobiotic induction, CncC induction or detoxification tissue enrichment) is not reliably diagnostic of detoxification function: not all detoxification genes are likely unstable over the 40–50 m.y. period analysed in this study, while essential E-class genes can sometimes be unstable (Feyereisen, 2020; Sztal et al., 2007); not all enzymes that detoxify a compound are induced by that compound (Willoughby et al., 2006); CncC regulates both detoxification and developmental gene sets (Deng and Kerppola, 2013; Misra et al., 2011); and ‘detoxification tissues’ have roles in non-detoxification processes, such as digestion, immunity, osmoregulation and energy metabolism (Beyenbach et al., 2010; Buchon et al., 2009; S. Li et al., 2019). S-class enzymes may also share evolutionary instability with X-class enzymes. Given this, any integrative method for detecting detoxification genes will produce some level of false positives and false negatives. It is also worth noting that the DS method as described here does not weight the importance of detoxification properties, even though, for example, detoxification tissue expression could arguably be more indicative of detoxification function than regulation by CncC. A more sophisticated, future version of this method should attempt this.

Loss of function alleles of *Cyp6a20* (DS = 3) result in aggressive behaviour in males, suggesting it may have a role in pheromone sensitivity (Dierick and Greenspan, 2006; Robin et al., 2007; L. Wang et al., 2008), although its enzymatic substrate is not known. Its possible misclassification as a detoxification candidate is due to its induction by both phenobarbital and methamphetamine, and its enrichment in the adult male midgut and the larval and adult female Malpighian tubules; it is also enriched in other tissues (Fig. S3), suggesting that it may have functions outside of the antennae. It is possible *Cyp6a20* encodes a bi-functional enzyme that belongs to both the S and X classes.

Indeed, the DS method might be limited more generally by the fact that some enzymes often fall into multiple functional classes simultaneously, such as the glutathione S-transferase GST16 in the cotton bollworm *Helicoverpa armigera*, which has roles in both detoxification and larval development (Shabab et al., 2014). There is experimental evidence that some P450 genes in *D. melanogaster* belong to multiple functional classes. *Cyp6g2* (DS = 1) may be involved in juvenile hormone metabolism (Christesen et al., 2016), but also has the capacity to detoxify multiple classes of insecticides if expressed ectopically (Daborn et al., 2007; Denecke et al., 2017). *Cyp6g2* is a paralog of *Cyp6g1* (DS = 4), which encodes a detoxification enzyme with very broad substrate specificity (Battlay et al., 2018; 2016; Daborn et al., 2002; 2007; Fusetto et al., 2017; Joußen et al., 2008; J. Schmidt et al., 2010), and so *Cyp6g2* may have a pre-adapted detoxification capacity. *Cyp6a17* (DS = 3) may detoxify insecticides (Battlay et al., 2018; Duneau et al., 2018) but also regulates temperature preference (Chakraborty et al., 2018; Kang et al., 2011). From an evolutionary perspective, the transition between E-, S- and X-class functions must occur with some frequency, and so it is likely that at any given time a non-zero proportion of enzymes in a gene family have multiple functions.

It has been previously noted that P450s harbouring polymorphic structural variation (SV; duplications, deletions or rearrangements) in *D. melanogaster* tend to belong to clades that are evolutionarily unstable between species (Good et al., 2014). We note that many high DS P450s with known detoxification functions identified in this study harbour SVs in the DGRP and the DSPR panels (Chakraborty et al., 2019; Good et al., 2014), including *Cyp6g1* (DS = 4)*, Cyp12d1* (DS = 4)*, Cyp28d1* (DS = 3), *Cyp6a17* (DS = 3) and *Cyp6a23* (DS = 3), suggesting that within-species SV might be a feature of detoxification genes more generally, complimenting between-species instability.

Due to the abundance of genome- and transcriptome-wide datasets in *D. melanogaster*, there are a number of other promising gene families that could be analysed by the DS method. The GSTs, CCEs and UDP-glycosyltransferases (UGTs) are all well-known detoxification families in *Drosophila* and other insects (Ahn et al., 2012; Low et al., 2007; Oakeshott et al., 2005; Rane et al., 2019) that have yet to be systematically studied this way and likely contain many undiscovered detoxification genes. Other gene families that appear in the detoxification-related genome-wide datasets used here could also be analysed using this method to explore whether or not they have detoxification functions.

The DS method could also be used to detect detoxification genes in non-model insects, such as agricultural pests or vectors of human disease. In order to do this, the following data would need to be generated: multiple genome assemblies of closely related species (or multiple genomes from the same species, to detect possible SV); detoxification tissue-specific and whole-body transcriptomics; and xenobiotic induction transcriptomic datasets. Transgenic manipulation of CncC and other detoxification-response factors has been described in multiple non-model species (reviewed in Wilding 2018). Utilizing these datasets, the DS method would likely produce candidate genes that could be validated with RNAi or CRISPR-Cas9 mutagenesis and lead to a greater understanding of detoxification in insects more generally.

### 4.4. Integration of EcKL and P450 PheWAS associations and candidate detoxification genes

Phenome-wide association studies (PheWAS) are powerful approaches to detect previously unknown associations between genes and phenotypes (Denny et al., 2010), and are particularly useful to study detoxification in *D. melanogaster*, as a large number of toxic stress phenotypes have been previously determined in this species. This study has uncovered a number of phenotypic associations with P450s and EcKLs that make intriguing candidates for future study.

In the P450 family, the association between a *Cyp4s3* (DS = 3) haplotype and developmental caffeine survival deserves particular attention, especially because the most strongly associated variant in the haplotype is a non-synonymous SNP. Of note, *Cyp4s3* is not known to be induced by exposure to caffeine in adults or larvae (Coelho et al., 2015; Willoughby et al., 2006), which is perhaps why it has yet to be studied in relation to caffeine detoxification. Caffeine is thought to be detoxified by multiple P450s in *D. melanogaster*, raising the possibility that *Cyp4s3* is involved in this process (Coelho et al., 2015).

In the EcKL family, *CG33301* (Dro26-2, DS = 3) has unique genomic variation associated with distinct phenotypic responses to three chemically disparate toxins: ethanol (adult tolerance; associated with a T4I amino acid substitution), imidacloprid (larval activity) and methylmercury (developmental tolerance). This suggests that either *CG33301* has a very broad substrate specificity or is involved in a stress-response process unrelated to direct detoxification. Its paralog, *CG16898* (Dro26-1, DS = 3), is also associated with methylmercury survival, suggesting the Dro26 clade as a whole may have toxic stress-related functions. *CG6908* (Dro23-0, DS = 4) transcript levels in male and female flies are strongly associated with both chlorantraniliprole and malathion survival. This may be due to its strong (66.5-fold) induction by CncC, which responds to oxidative stress and regulates a core subset of genes transcriptionally associated with resistance to both insecticides (Battlay et al., 2018; L. Green et al., 2019). Because CncC positively regulates the transcription of hundreds of genes, and its constitutive up-regulation in a subset of DGRP lines has produced a co-transcriptional module (L. Green et al., 2019), it is possible *CG6908* is not functionally connected to chlorantraniliprole and malathion detoxification and the associations may simply be due to its co-regulation with *Cyp12d1*, which transgenic experiments suggest directly affects resistance to these insecticides (Battlay et al., 2018; L. Green et al., 2019). Regardless, *CG6908*’s high detoxification score suggests the gene is involved in detoxification in some capacity and requires further study. *CG10560* (Dro17-3, DS = 4) expression is also associated with chlorantraniliprole survival, but it is also strongly induced by CncC (16.2-fold), suggesting this may also be a non-casual association. *CG11878* (Dro1-5) transcript levels in female flies are associated with ethanol tolerance in male flies, which might suggest the association is spurious, but this gene is also up-regulated in response to ethanol exposure in male flies (Morozova et al., 2006), which tends to strengthen the link.

### 4.5. Functional characterisation of EcKLs

Prior to the current study, very few EcKLs in *D. melanogaster* have been functionally characterised. *JhI-26* (Dro46-0, DS = 3) is positively regulated by juvenile hormone (Dubrovsky et al., 2000) and forms a putatively causal link between both the increased juvenile hormone titre and increased expression of the male accessory gland protein CG10433 found during *Wolbachia* infection (Liu et al., 2014). Overexpression of *JhI-26* in the testes of male flies results in paternal-effect lethality and a reduction in the mating receptivity of females with which they have mated, suggesting it plays a role in *Wolbachia*-mediated cytoplasmic incompatibility (Liu et al., 2014). Curiously, *JhI-26*’s DS value determined in this study suggests it may also have a role in detoxification.

*CHKov1* (Dro18-2, DS = 2) and *CHKov2* (Dro19-0, DS = 2) have also been studied previously; a *CHKov1* TE-insertion allele has undergone a selective sweep in the last 200 years, which was putatively linked to organophosphate insecticide resistance by Aminetzach et al. (2005). However, this allele—and another containing duplications of *CHKov1* and *CHKov2*—also clearly confer strong resistance to the sigma virus (Magwire et al., 2012; 2011). While Aminetzach et al. (2005) provide evidence the original TE allele confers resistance to the organophosphate insecticide azinphos-methyl using a single backcrossed line, no association between the *CHKov1*-TE allele and azinphos-methyl resistance was found in a recent genome-wide association study using a large panel of inbred lines (Battlay et al., 2016). Our results here suggest neither *CHKov1* nor *CHKov2* is a detoxification gene, although the paralog of *CHKov1*, *CG10562* (Dro18-1), is a detoxification candidate.

Although we have presented preliminary evidence in this study for the hypothesis that the EcKLs are a detoxification family, it is likely that some genes in the family have E-class functions, such as in ecdysteroid metabolism (Sonobe and Ito, 2009). Genes with low detoxification scores (0 or 1) may have important developmental or physiological functions, as seen in the P450s (Table 2), in which case up to 12 EcKLs (Fig. 8) could be considered candidate E-class genes. Of these, *CG31102* (Dro9-0), *CG31099* (Dro14-0) and *CG9497* (Dro31-0) show developmental lethality upon putative knockdown in this study (Fig. S9) and are the strongest candidates for E-class EcKLs in *D. melanogaster*. However, it is important to note that some RNAi constructs fail to knock down mRNA transcript levels sufficiently to observe a phenotype, while others can produce substantial off-target effects (Heigwer et al., 2018); given this, phenotypes observed with only a single RNAi construct should be treated as tentative until further characterisation is conducted, and lack of a phenotype cannot be considered evidence for a lack of developmental essentiality. With this in mind, we assert that little can be concluded from the RNAi knockdown data presented in this study without follow-up work. Other good E-class candidate EcKLs are *CG7135* (Dro32-0), *CG1561* (Dro37-0), *CG14314* (Dro40-0) and *CG5644* (Dro42-0), which all have detoxification scores of 0 and are highly conserved in the *Drosophila* genus.

*D. melanogaster* is associated with a powerful collection of genetic tools for the functional characterisation of detoxification genes and provides an ideal model for studying insect detoxification and the function of EcKLs in a general sense (Scott and Buchon, 2019). However, since detoxification can help define ecological niches for some insects and therefore be quite specific for certain toxins, it will be necessary to extend the functional characterisation of EcKLs to non-*Drosophila* species.

If EcKLs are responsible for detoxicative phosphorylation observed in insects, this would imply their X-class substrates may include phenols, steroids and glucosides (Table 1). Phytoecdysteroids and mycoecdysteroids are present in various plant and fungal species as anti-insect secondary metabolites (Dinan, 2001; Kovganko, 1999), and some are likely detoxified by phosphorylation (Rharrabe et al., 2007). Other hydroxylated toxins that could be X-class EcKL substrates include withanolides, cardenolides, cucurbitacins and flavonoids (Agrawal et al., 2012; Dinan et al., 1997; Glotter, 2011; S. Wang et al., 2017). X-class EcKLs may also phosphorylate the products of phase I hydroxylation reactions or phase II glucosidation reactions, as is speculated to occur in locusts and moths (Boeckler et al., 2016; Olsen et al., 2015; 2014).

### 5. Summary

In this study, we have shown that it is possible to use the abundant genomic and transcriptomic resources in the *Drosophila* genus to test functional hypotheses about a poorly understood gene family, the EcKLs. We have also highlighted that there is much more to discover about the functions of the P450 family in insects, particularly with respect to xenobiotic metabolism. We hope that this work can be used as a springboard for further characterisation of both gene families, and that it might inspire similar work in other insect taxa.

## Author contributions

J.L.S. and P.B.: Writing the manuscript

J.L.S., R.S.G-S., P.B. and C.R.: Editing the manuscript

J.L.S. and R.S.G-S.: Laboratory work

J.L.S. and P.B.: Analysis of data

C.R.: Funding and overall project design

## Acknowledgements

We thank Rob Good and Jin Kee for conversations and analyses that helped motivate this research, as well as Dr Lars Jermiin for his supervisory wisdom and financial support, and Melanie Stewart for feedback on the manuscript. We would also like to thank Pontus Leblanc for assisting with the RNAi knockdown crosses. This research was partially supported by ARC grant DP985013 and University of Melbourne support to C.R.

---

**Figure S1:**
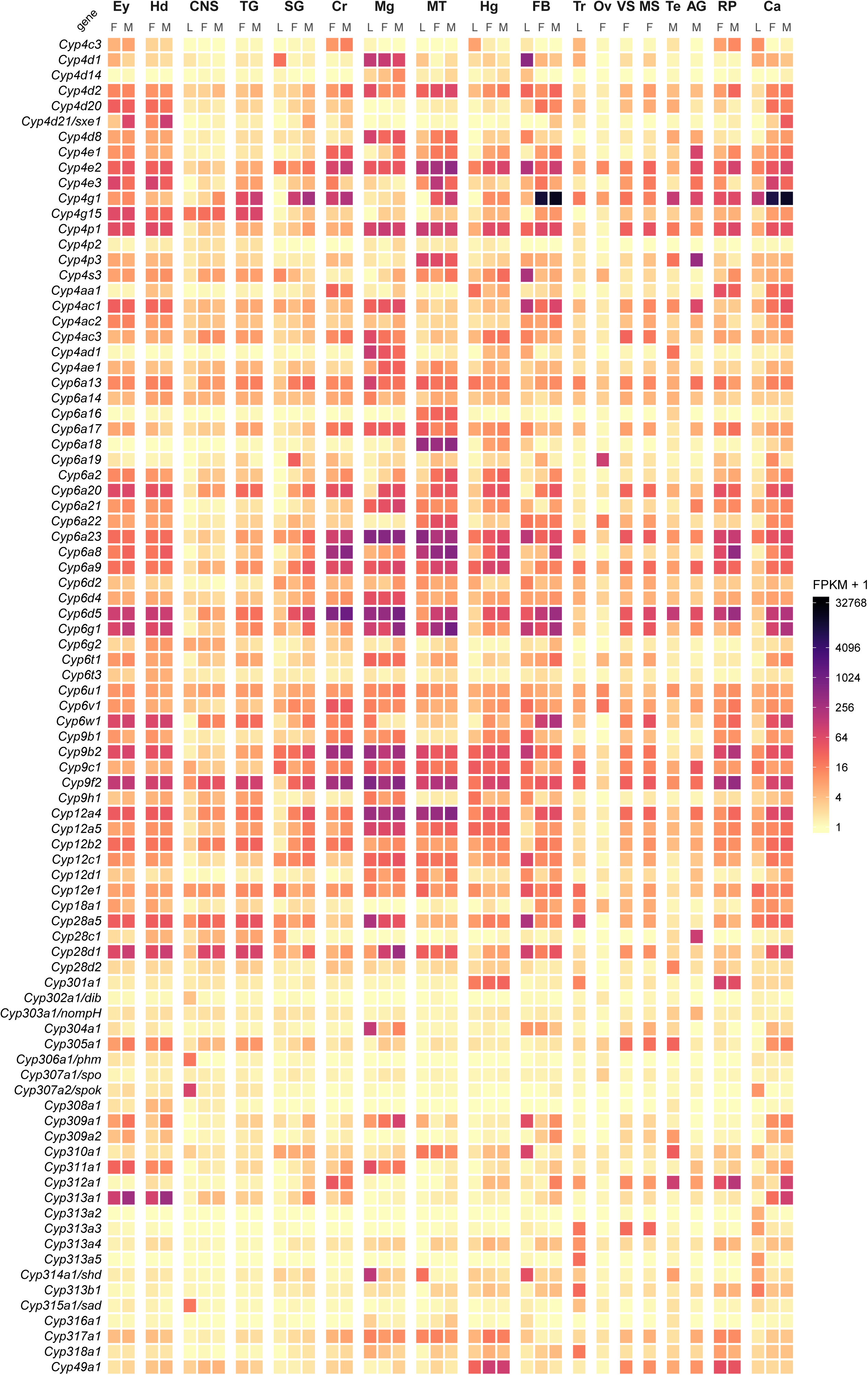
Heatmap of absolute expression (FPKM) of P450 genes in *Drosophila melanogaster* across 18 tissues and three life stages. L, 3^rd^ instar larva; M, adult male; F, adult female. Ey, eye; Hd, head; CNS, central nervous system; TG, Thoracicoabdominal ganglion; SG, salivary gland; Cr, crop; Mg, midgut; MT, Malpighian tubules; Hg, hindgut; FB, fat body; Tr, trachea; Ov, ovary; VS, virgin spermatheca; MS, mated spermatheca; Te, testis; AG, accessory gland; RP, rectal pad; Ca, carcass.

**Figure S2:**
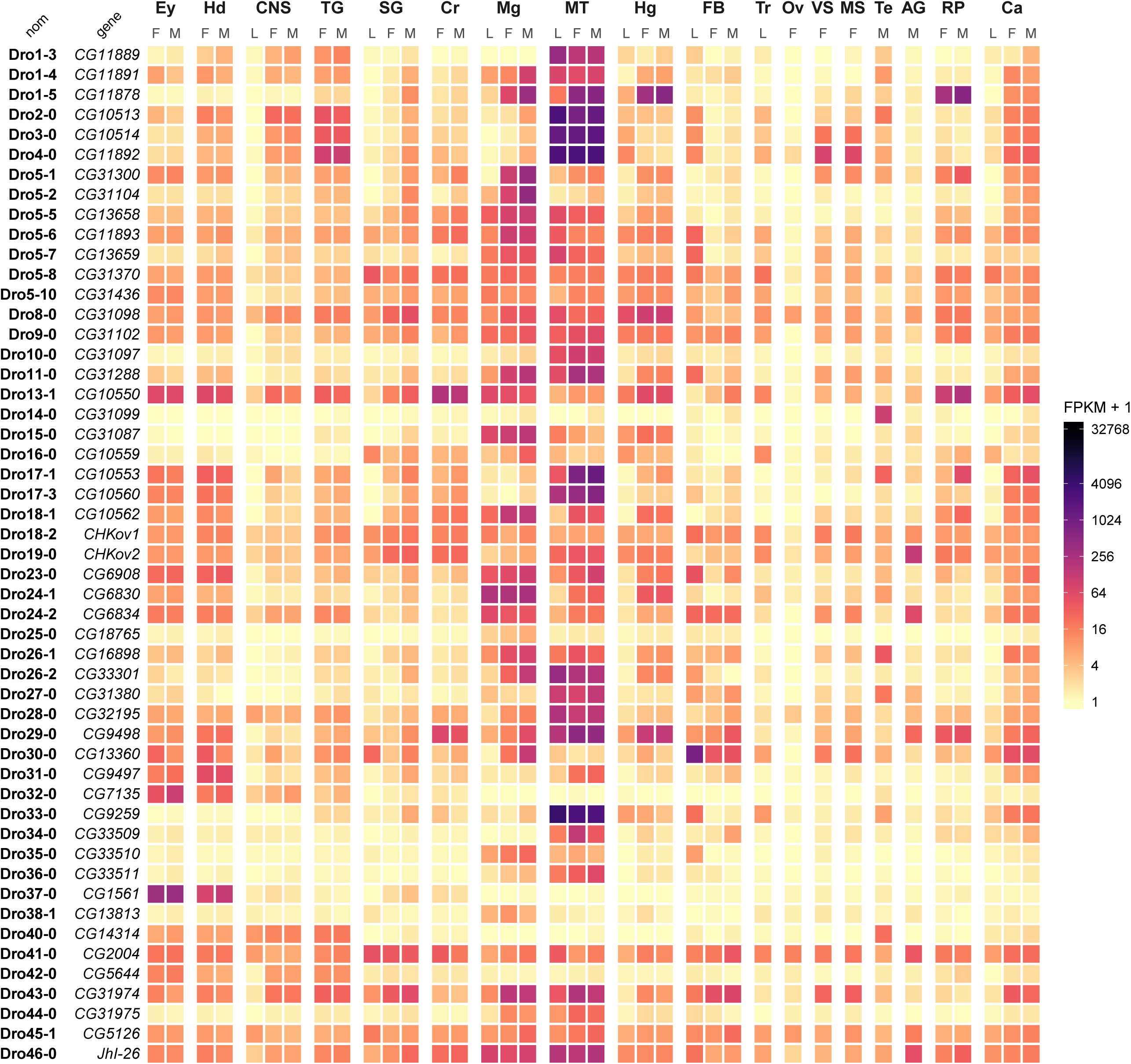
Heatmap of absolute expression (FPKM) of EcKL genes in *Drosophila melanogaster* across 18 tissues and three life stages. L, 3^rd^ instar larva; M, adult male; F, adult female. Ey, eye; Hd, head; CNS, central nervous system; TG, Thoracicoabdominal ganglion; SG, salivary gland; Cr, crop; Mg, midgut; MT, Malpighian tubules; Hg, hindgut; FB, fat body; Tr, trachea; Ov, ovary; VS, virgin spermatheca; MS, mated spermatheca; Te, testis; AG, accessory gland; RP, rectal pad; Ca, carcass.

**Figure S3:**
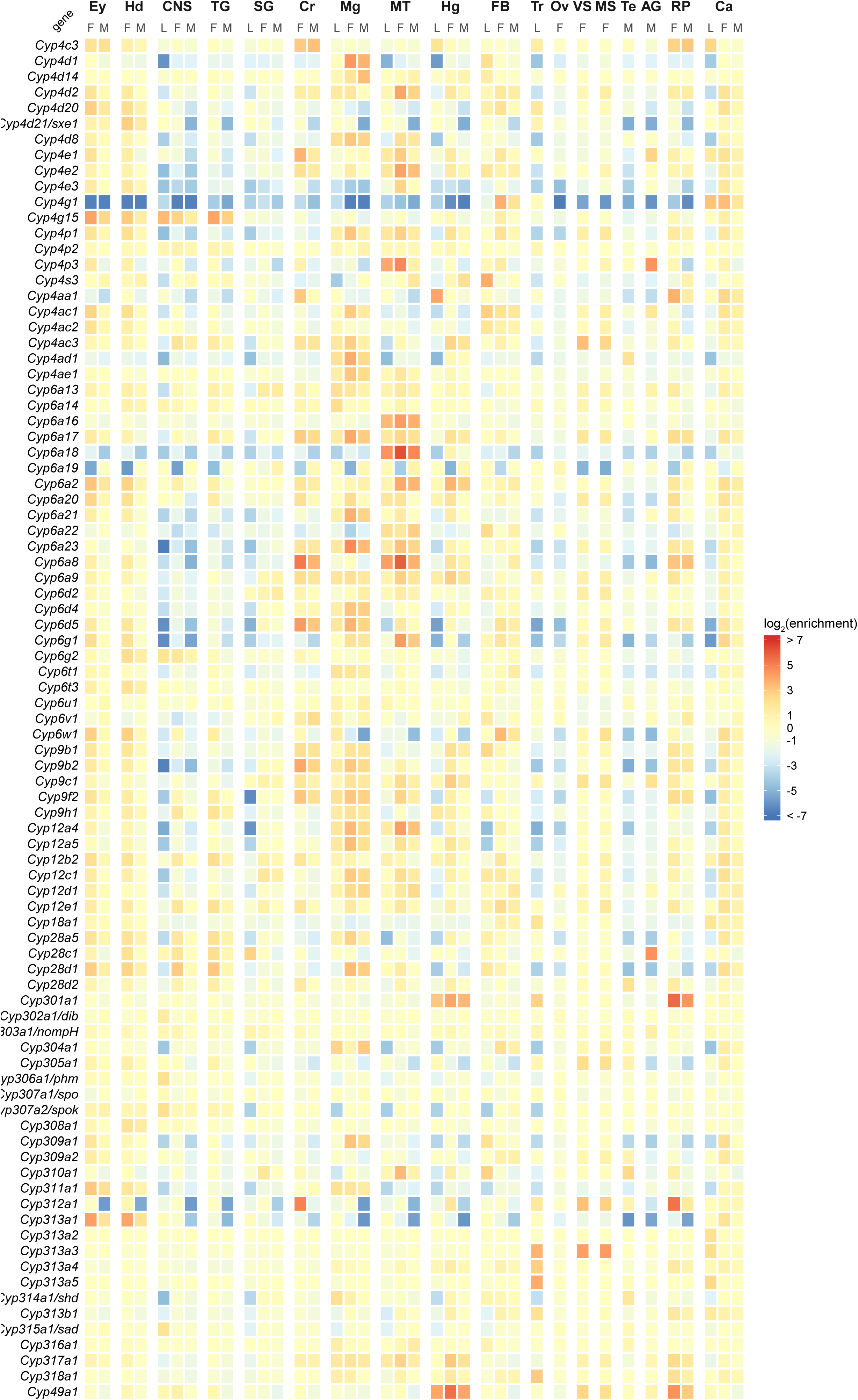
Heatmap of log_2_(enrichment) of P450 genes in *Drosophila melanogaster* across 18 tissues and three life stages. L, 3^rd^ instar larva; M, adult male; F, adult female. Ey, eye; Hd, head; CNS, central nervous system; TG, Thoracicoabdominal ganglion; SG, salivary gland; Cr, crop; Mg, midgut; MT, Malpighian tubules; Hg, hindgut; FB, fat body; Tr, trachea; Ov, ovary; VS, virgin spermatheca; MS, mated spermatheca; Te, testis; AG, accessory gland; RP, rectal pad; Ca, carcass.

**Figure S4:**
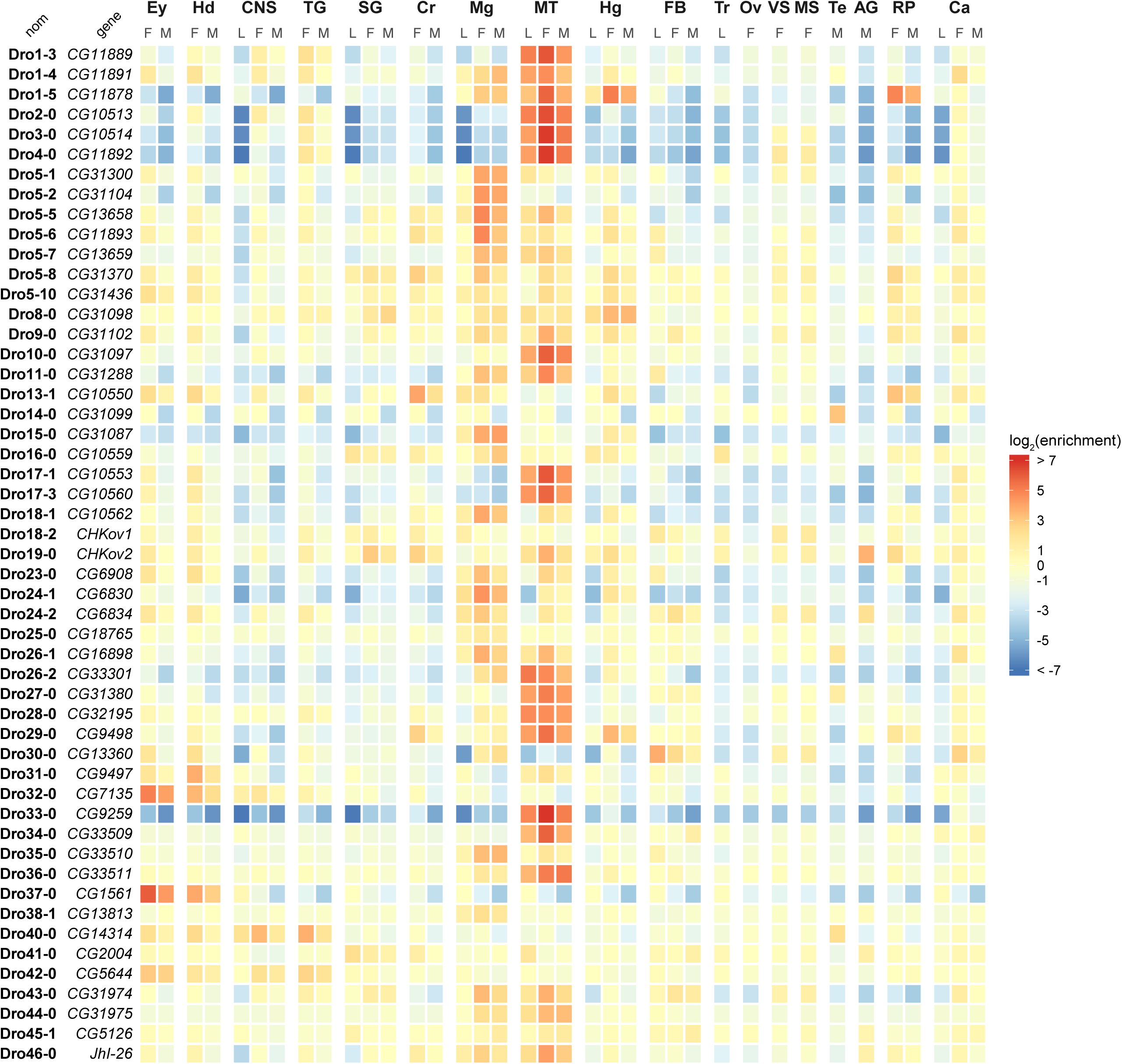
Heatmap of log_2_(enrichment) of EcKL genes in *Drosophila melanogaster* across 18 tissues and three life stages. L, 3^rd^ instar larva; M, adult male; F, adult female. Ey, eye; Hd, head; CNS, central nervous system; TG, Thoracicoabdominal ganglion; SG, salivary gland; Cr, crop; Mg, midgut; MT, Malpighian tubules; Hg, hindgut; FB, fat body; Tr, trachea; Ov, ovary; VS, virgin spermatheca; MS, mated spermatheca; Te, testis; AG, accessory gland; RP, rectal pad; Ca, carcass.

**Figure S5:**
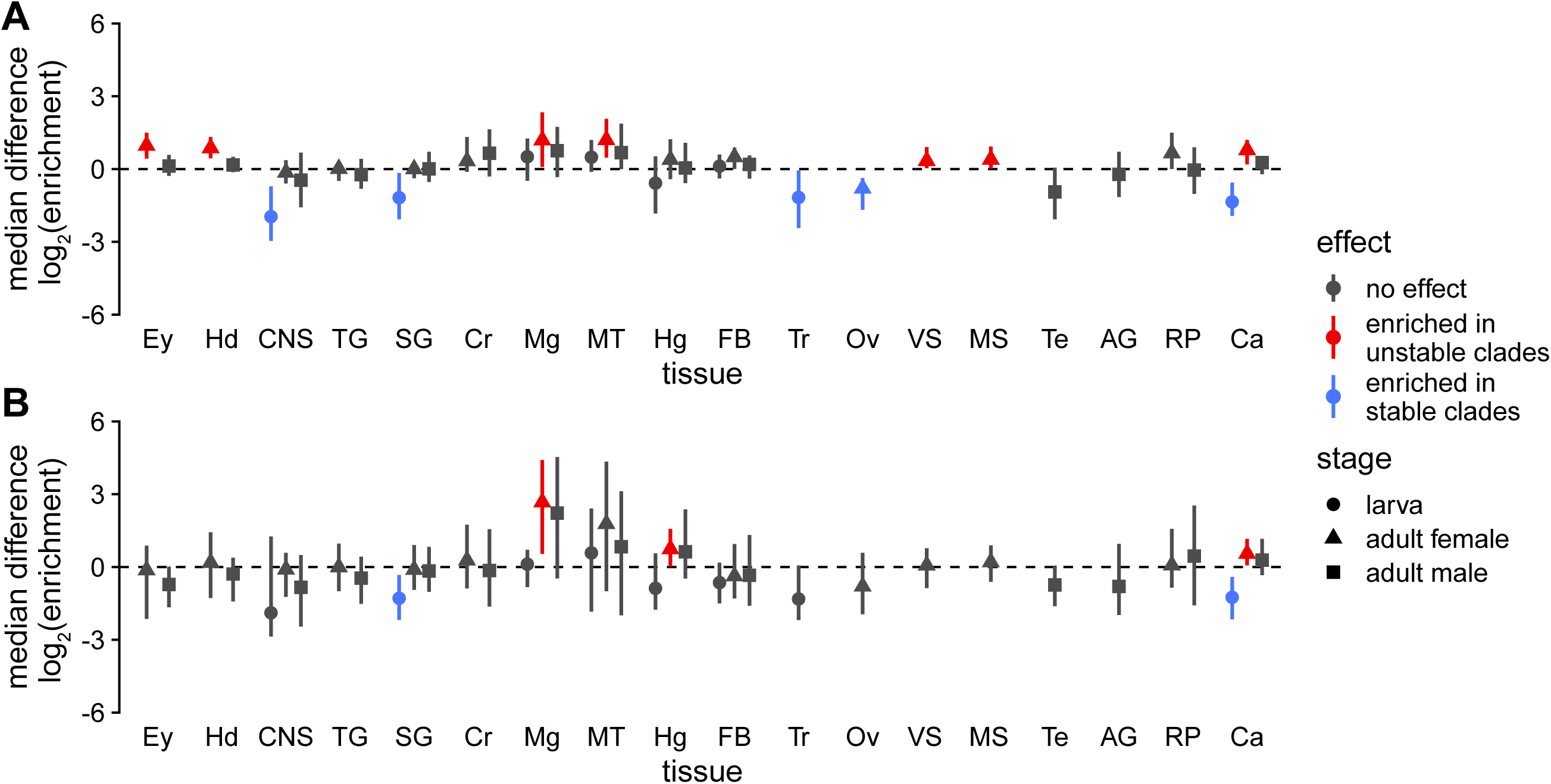
Median log_2_(enrichment) differences between unstable and stable genes in the P450 (A) and EcKL (B) gene families for specific tissues and life stages. Error bars are 95% confidence intervals; effects were considered significant if the interval did not overlap with 0. Ey, eye; Hd, head; CNS, central nervous system; TG, Thoracicoabdominal ganglion; SG, salivary gland; Cr, crop; Mg, midgut; MT, Malpighian tubules; Hg, hindgut; FB, fat body; Tr, trachea; Ov, ovary; VS, virgin spermatheca; MS, mated spermatheca; Te, testis; AG, accessory gland; RP, rectal pad; Ca, carcass.

**Figure S6:**
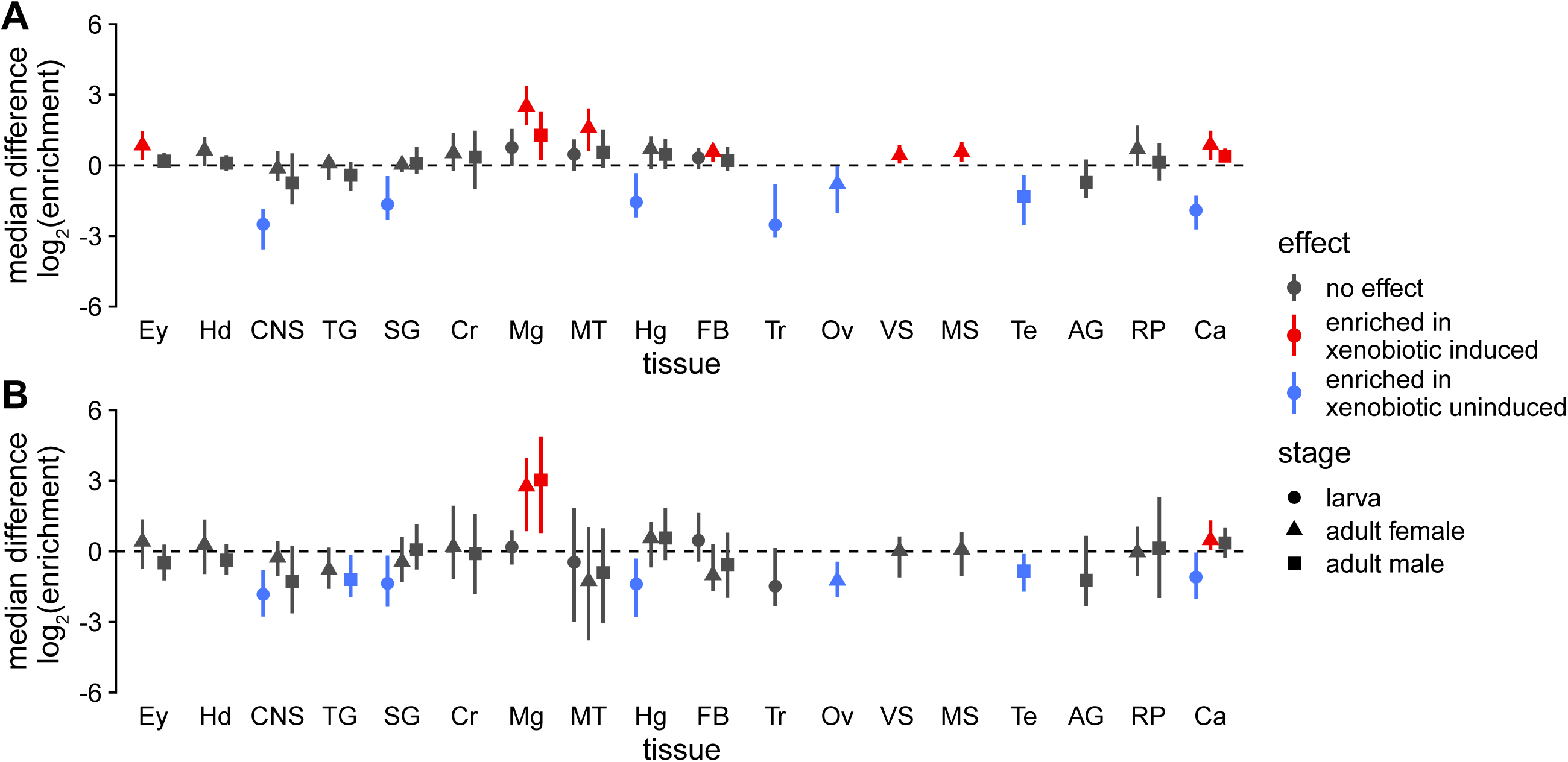
Median log_2_(enrichment) differences between xenobiotically induced and uninduced genes in the P450 (A) and EcKL (B) gene families for specific tissues and life stages. Error bars are 95% confidence intervals; effects were considered significant if the interval did not overlap with 0. Ey, eye; Hd, head; CNS, central nervous system; TG, Thoracicoabdominal ganglion; SG, salivary gland; Cr, crop; Mg, midgut; MT, Malpighian tubules; Hg, hindgut; FB, fat body; Tr, trachea; Ov, ovary; VS, virgin spermatheca; MS, mated spermatheca; Te, testis; AG, accessory gland; RP, rectal pad; Ca, carcass.

**Figure S7:**
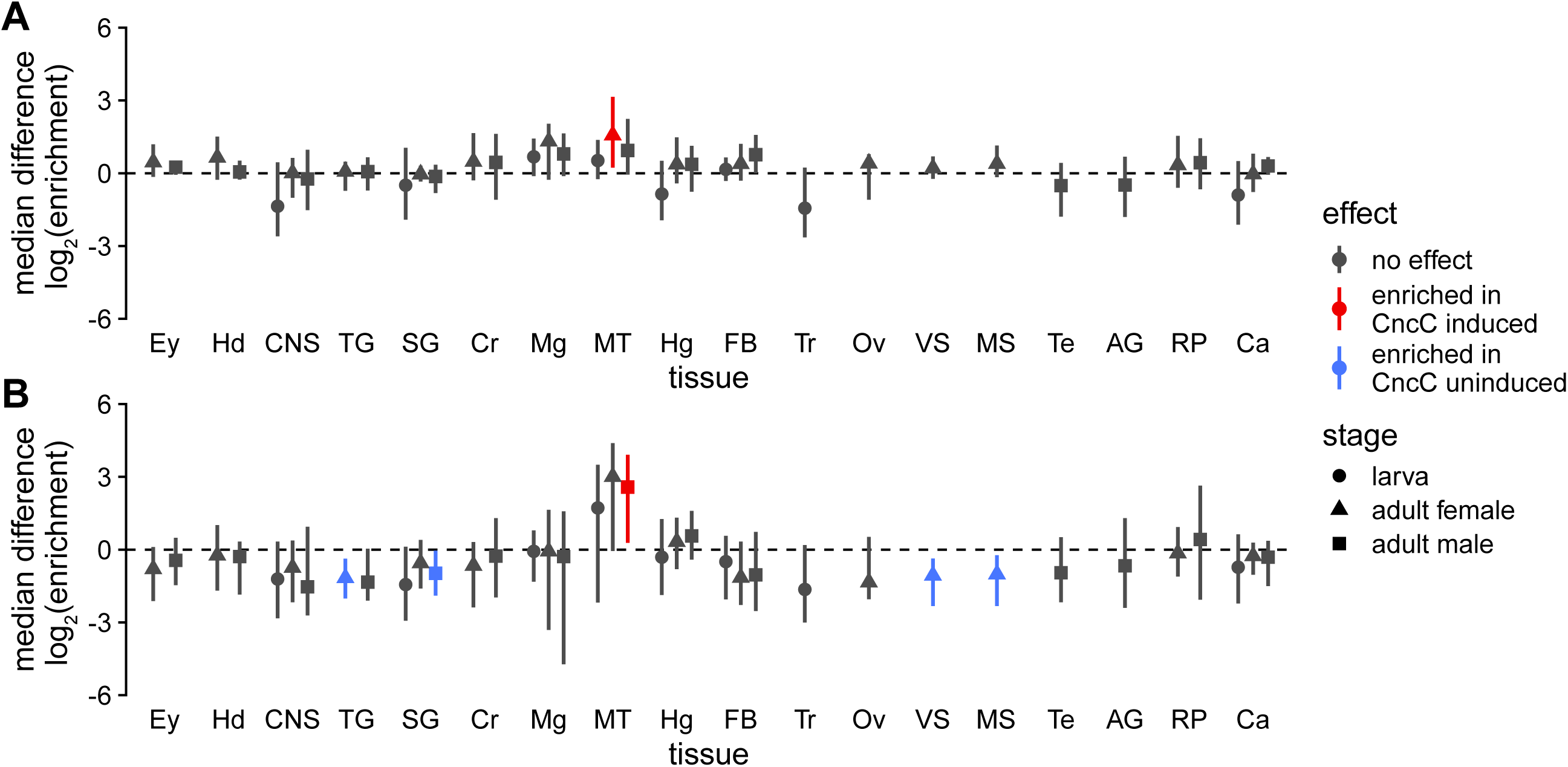
Median log_2_(enrichment) differences between CncC induced and uninduced genes in the P450 (A) and EcKL (B) gene families for specific tissues and life stages. Error bars are 95% confidence intervals; effects were considered significant if the interval did not overlap with 0. Ey, eye; Hd, head; CNS, central nervous system; TG, Thoracicoabdominal ganglion; SG, salivary gland; Cr, crop; Mg, midgut; MT, Malpighian tubules; Hg, hindgut; FB, fat body; Tr, trachea; Ov, ovary; VS, virgin spermatheca; MS, mated spermatheca; Te, testis; AG, accessory gland; RP, rectal pad; Ca, carcass.

**Figure S8:**
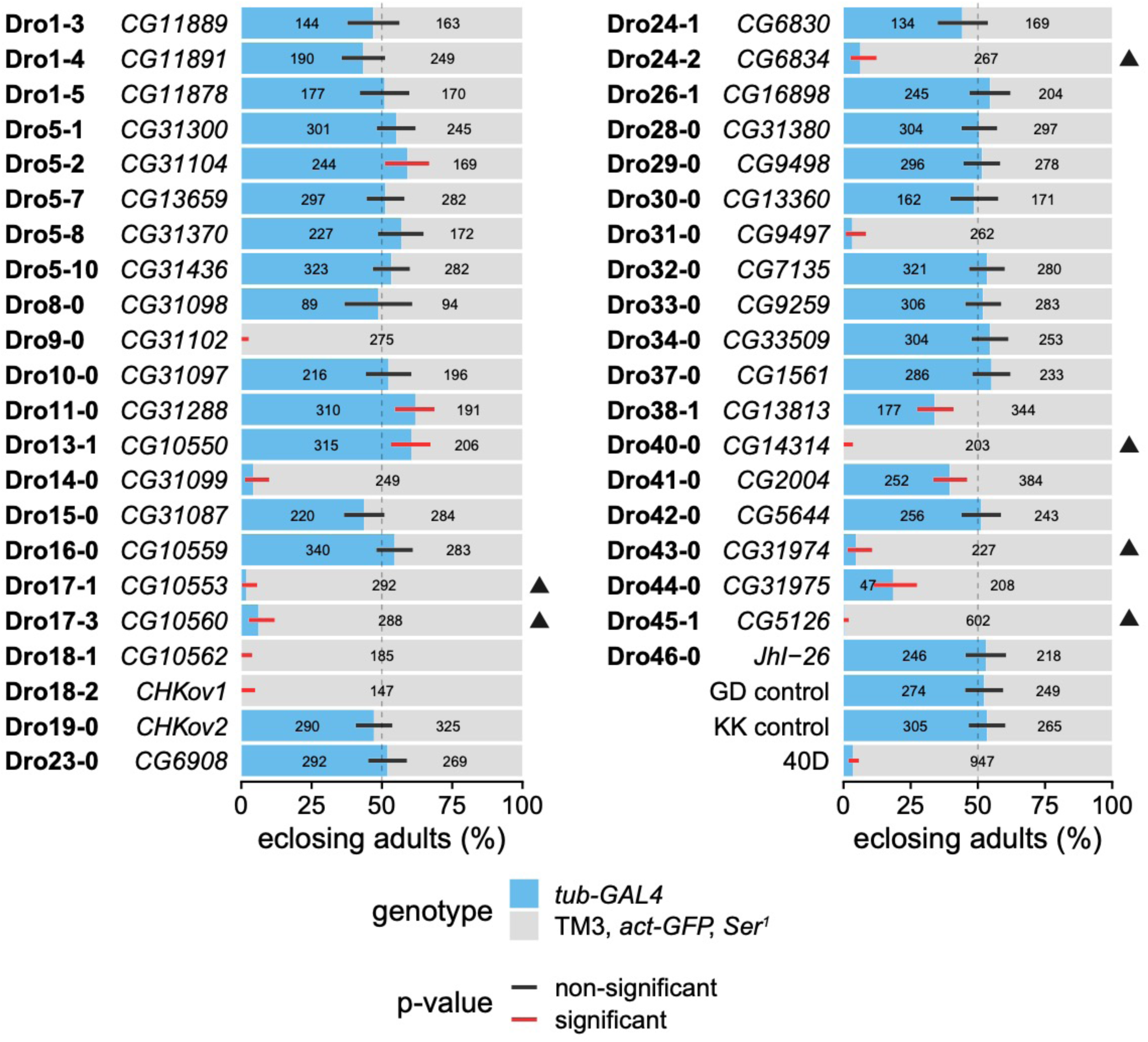
RNAi knockdown of 41 EcKL genes in *Drosophila melanogaster* and three control crosses, scoring offspring genotypes. Error bars are 99.89% confidence intervals (95% confidence interval adjusted for 44 tests) for the proportion of *tub-GAL4* individuals; black and red bars indicate non-significant or significant deviations, respectively, from expected genotypic ratios after correction for multiple tests. For the *CG31098* cross, the UAS-dsRNA construct was over a CyO balancer, so only non-CyO individuals have been scored. For the *CHKov1* cross, the UAS-dsRNA construct was on the X-chromosome, so only female individuals (inheriting the driver and responder) have been scored. The number of eclosed adults of each genotype is indicated by the number within each bar (numbers less than 40 are not shown). Results from KK library dsRNA lines with annotated insertions of the hairpin are indicated with black triangles; 40D is a line containing a UAS-only insertion at the annotated position for the KK library and acts as a positive control for *tiptop*-related phenotypes. dsRNA VDRC line IDs can be found in Table S4.

