## Supplementary material for "Identifying candidate detoxification genes in the ecdysteroid kinase-like (EcKL) and cytochrome P450 gene families in *Drosophila melanogaster* by integrating evolutionary and transcriptomic data": Figures S1-8

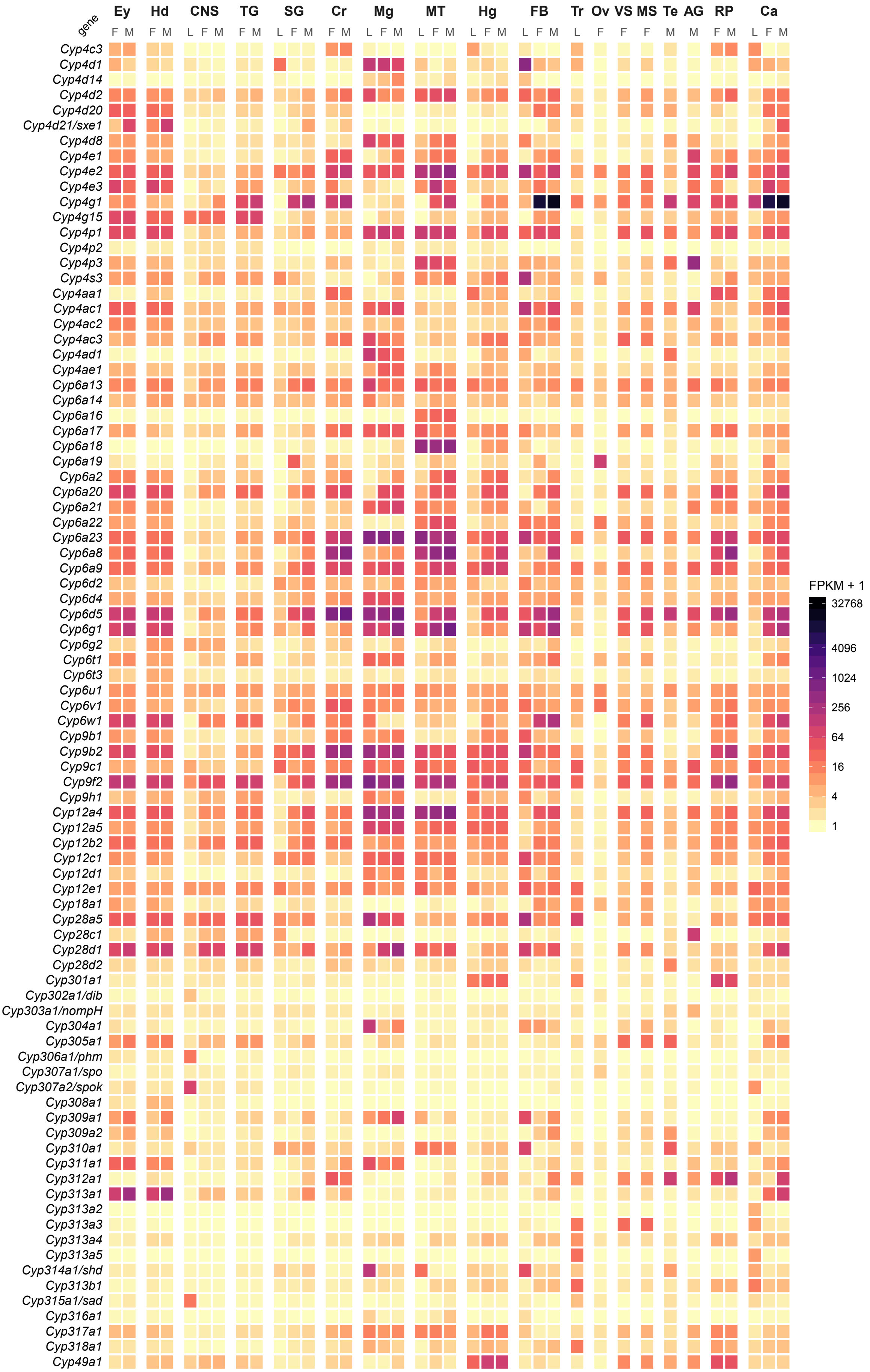

gene
Ey
Hd
CNS
TG
SG
Cr
Mg
MT
Hg
FB
Tr
Ov
VS
MS
Te
AG
RP
Ca
F M F M L F M F M L F M F M L F M L F M L F M L F M L F F F M M F M L F M
Cyp4c3
Cyp4d1
Cyp4d14
Cyp4d2
Cyp4d20
Cyp4d21/sxe1
Cyp4d8
Cyp4e1
Cyp4e2
Cyp4e3
Cyp4g1
Cyp4g15
Cyp4p1
Cyp4p2
Cyp4p3
Cyp4s3
Cyp4aa1
Cyp4ac1
Cyp4ac2
Cyp4ac3
Cyp4ad1
Cyp4ae1
Cyp6a13
Cyp6a14
Cyp6a16
Cyp6a17
Cyp6a18
Cyp6a19
Cyp6a2
Cyp6a20
Cyp6a21
Cyp6a22
Cyp6a23
Cyp6a8
Cyp6a9
Cyp6d2
Cyp6d4
Cyp6d5
Cyp6g1
Cyp6g2
Cyp6t1
Cyp6t3
Cyp6u1
Cyp6v1
Cyp6w1
Cyp9b1
Cyp9b2
Cyp9c1
Cyp9f2
Cyp9h1
Cyp12a4
Cyp12a5
Cyp12b2
Cyp12c1
Cyp12d1
Cyp12e1
Cyp18a1
Cyp28a5
Cyp28c1
Cyp28d1
Cyp28d2
Cyp301a1
Cyp302a1/dib
Cyp303a1/nompH
Cyp304a1
Cyp305a1
Cyp306a1/phm
Cyp307a1/spo
Cyp307a2/spok
Cyp308a1
Cyp309a1
Cyp309a2
Cyp310a1
Cyp311a1
Cyp312a1
Cyp313a1
Cyp313a2
Cyp313a3
Cyp313a4
Cyp313a5
Cyp314a1/shd
Cyp313b1
Cyp315a1/sad
Cyp316a1
Cyp317a1
Cyp318a1
Cyp49a1
FPKM + 1
32768
4096
1024
256
64
16
4
1

**Figure S1:** Heatmap of absolute expression (FPKM) of P450 genes in *Drosophila melanogaster* across 18 tissues and three life stages. L, 3rd instar larva; M, adult male; F, adult female. Ey, eye; Hd, head; CNS, central nervous system; TG, Thoracicoabdominal ganglion; SG, salivary gland; Cr, crop; Mg, midgut; MT, Malpighian tubules; Hg, hindgut; FB, fat body; Tr, trachea; Ov, ovary; VS, virgin spermatheca; MS, mated spermatheca; Te, testis; AG, accessory gland; RP, rectal pad; Ca, carcass.

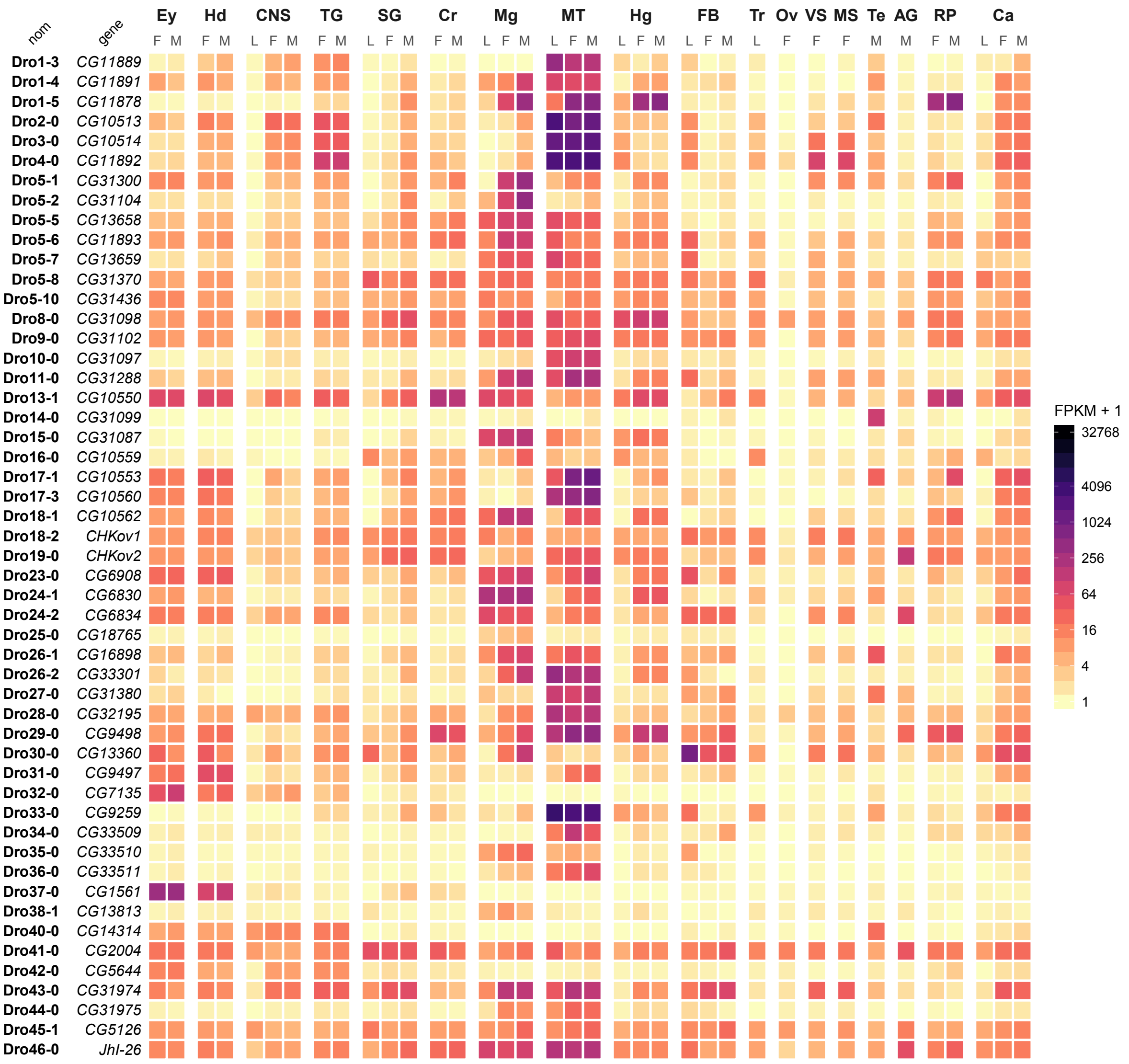


**Figure S2:** Heatmap of absolute expression (FPKM) of EcKL genes in *Drosophila melanogaster* across 18 tissues and three life stages. L, 3rd instar larva; M, adult male; F, adult female. Ey, eye; Hd, head; CNS, central nervous system; TG, Thoracicoabdominal ganglion; SG, salivary gland; Cr, crop; Mg, midgut; MT, Malpighian tubules; Hg, hindgut; FB, fat body; Tr, trachea; Ov, ovary; VS, virgin spermatheca; MS, mated spermatheca; Te, testis; AG, accessory gland; RP, rectal pad; Ca, carcass.

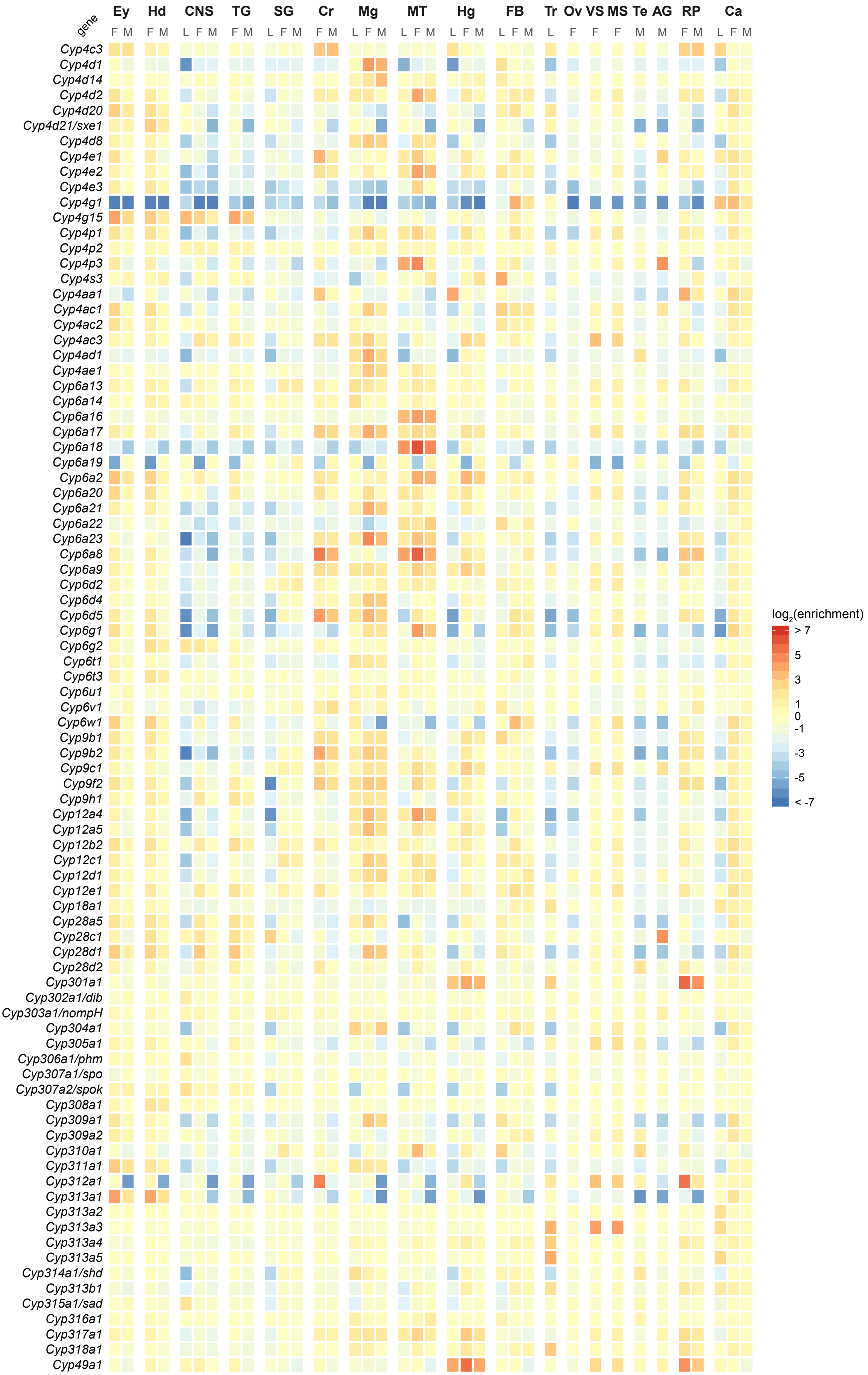
Ey
Hd
CNS
TG
SG
Cr
Mg
MT
Hg
FB
Tr
Ov
VS
MS
Te
AG
RP
Ca
gene
F M
F M
L F M
F M
L F M
F M
L F M
L F M
L F M
L F M
L
F
F
F
M
M
F M
L F M
Cyp4c3
Cyp4d1
Cyp4d14
Cyp4d2
Cyp4d20
Cyp4d21/sxe1
Cyp4d8
Cyp4e1
Cyp4e2
Cyp4e3
Cyp4g1
Cyp4g15
Cyp4p1
Cyp4p2
Cyp4p3
Cyp4s3
Cyp4aa1
Cyp4ac1
Cyp4ac2
Cyp4ac3
Cyp4ad1
Cyp4ae1
Cyp6a13
Cyp6a14
Cyp6a16
Cyp6a17
Cyp6a18
Cyp6a19
Cyp6a2
Cyp6a20
Cyp6a21
Cyp6a22
Cyp6a23
Cyp6a8
Cyp6a9
Cyp6d2
Cyp6d4
Cyp6d5
Cyp6g1
Cyp6g2
Cyp6t1
Cyp6t3
Cyp6u1
Cyp6v1
Cyp6w1
Cyp9b1
Cyp9b2
Cyp9c1
Cyp9f2
Cyp9h1
Cyp12a4
Cyp12a5
Cyp12b2
Cyp12c1
Cyp12d1
Cyp12e1
Cyp18a1
Cyp28a5
Cyp28c1
Cyp28d1
Cyp28d2
Cyp301a1
Cyp302a1/dib
Cyp303a1/nompH
Cyp304a1
Cyp305a1
Cyp306a1/phm
Cyp307a1/spo
Cyp307a2/spok
Cyp308a1
Cyp309a1
Cyp309a2
Cyp310a1
Cyp311a1
Cyp312a1
Cyp313a1
Cyp313a2
Cyp313a3
Cyp313a4
Cyp313a5
Cyp314a1/shd
Cyp313b1
Cyp315a1/sad
Cyp316a1
Cyp317a1
Cyp318a1
Cyp49a1
log2(enrichment)
> 7
5
3
1
0
-1
-3
-5
< -7

**Figure S3:** Heatmap of $log_2$(enrichment) of P450 genes in *Drosophila melanogaster* across 18 tissues and three life stages. L, $3^{rd}$ instar larva; M, adult male; F, adult female. Ey, eye; Hd, head; CNS, central nervous system; TG, Thoracicoabdominal ganglion; SG, salivary gland; Cr, crop; Mg, midgut; MT, Malpighian tubules; Hg, hindgut; FB, fat body; Tr, trachea; Ov, ovary; VS, virgin spermatheca; MS, mated spermatheca; Te, testis; AG, accessory gland; RP, rectal pad; Ca, carcass.

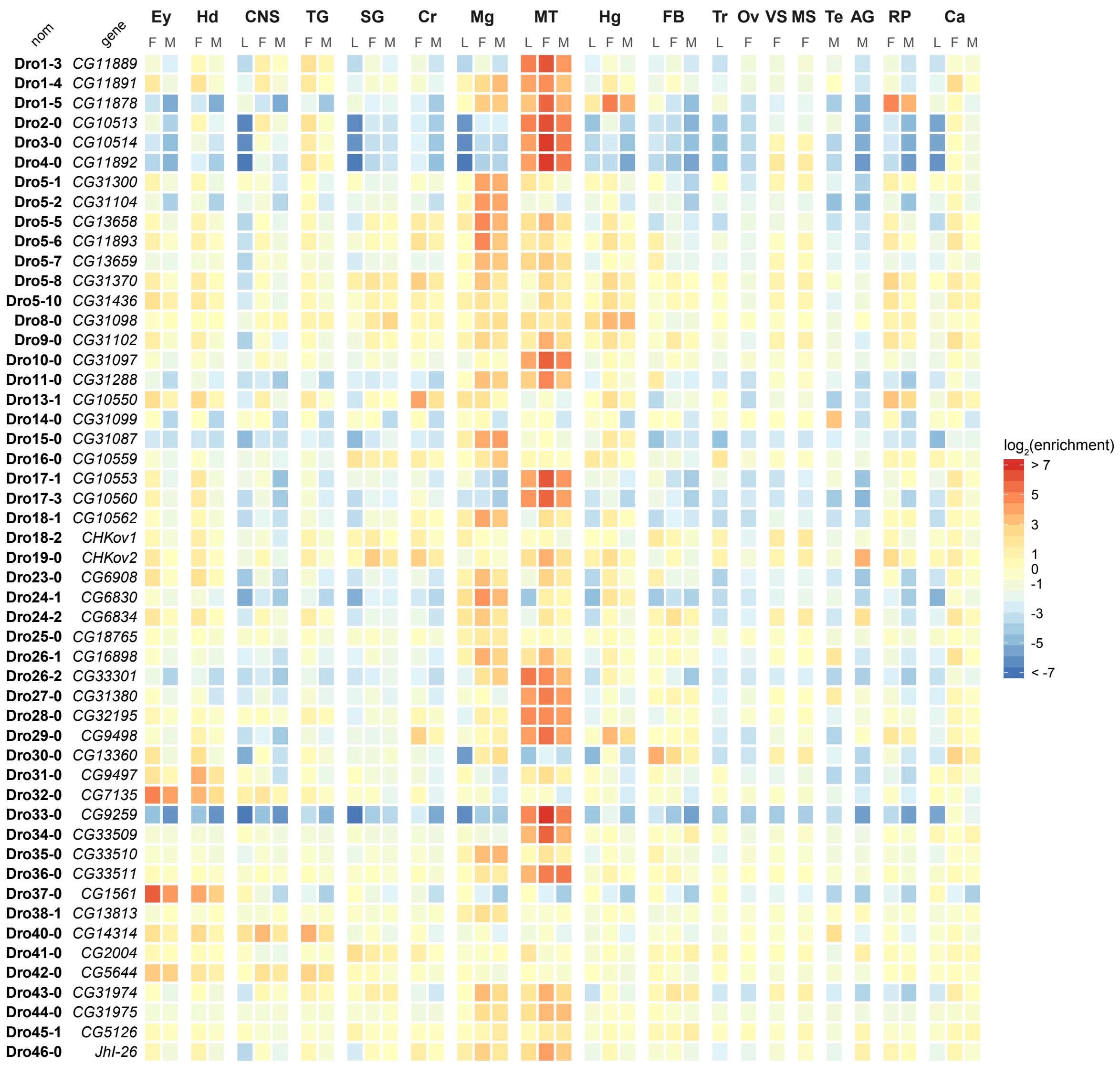


**Figure S4:** Heatmap of $log_2$(enrichment) of EcKL genes in *Drosophila melanogaster* across 18 tissues and three life stages. L, 3rd instar larva; M, adult male; F, adult female. Ey, eye; Hd, head; CNS, central nervous system; TG, Thoracicoabdominal ganglion; SG, salivary gland; Cr, crop; Mg, midgut; MT, Malpighian tubules; Hg, hindgut; FB, fat body; Tr, trachea; Ov, ovary; VS, virgin spermatheca; MS, mated spermatheca; Te, testis; AG, accessory gland; RP, rectal pad; Ca, carcass.

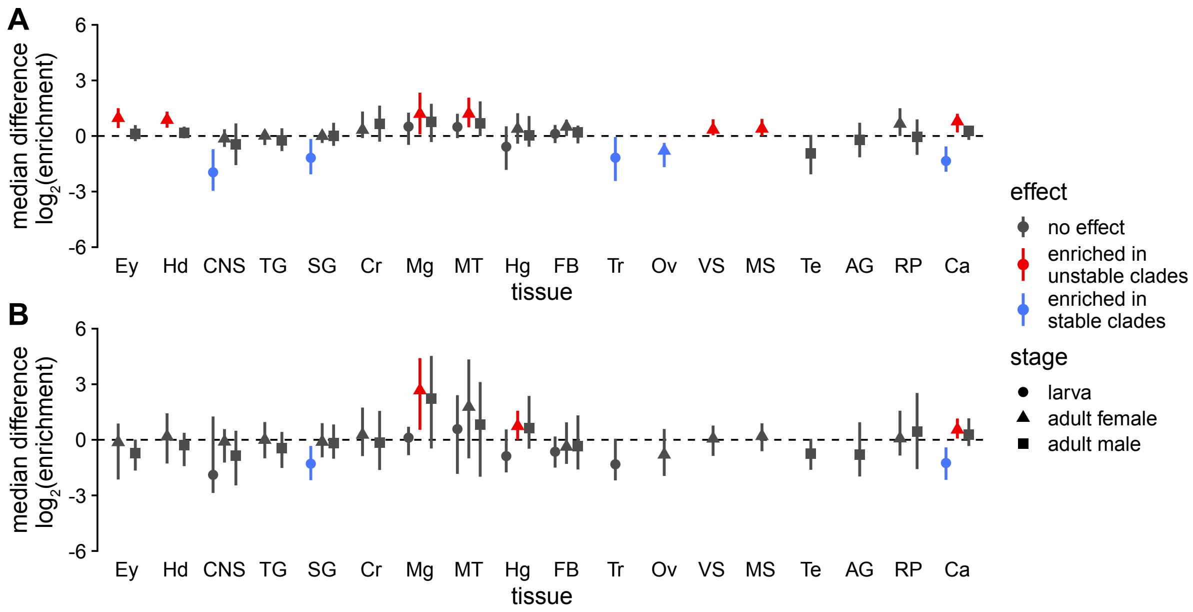


**Figure S5:** Median $\log_2$(enrichment) differences between unstable and stable genes in the P450 (A) and EcKL (B) gene families for specific tissues and life stages. Error bars are 95% confidence intervals; effects were considered significant if the interval did not overlap with 0. Ey, eye; Hd, head; CNS, central nervous system; TG, Thoracicoabdominal ganglion; SG, salivary gland; Cr, crop; Mg, midgut; MT, Malpighian tubules; Hg, hindgut; FB, fat body; Tr, trachea; Ov, ovary; VS, virgin spermatheca; MS, mated spermatheca; Te, testis; AG, accessory gland; RP, rectal pad; Ca, carcass.

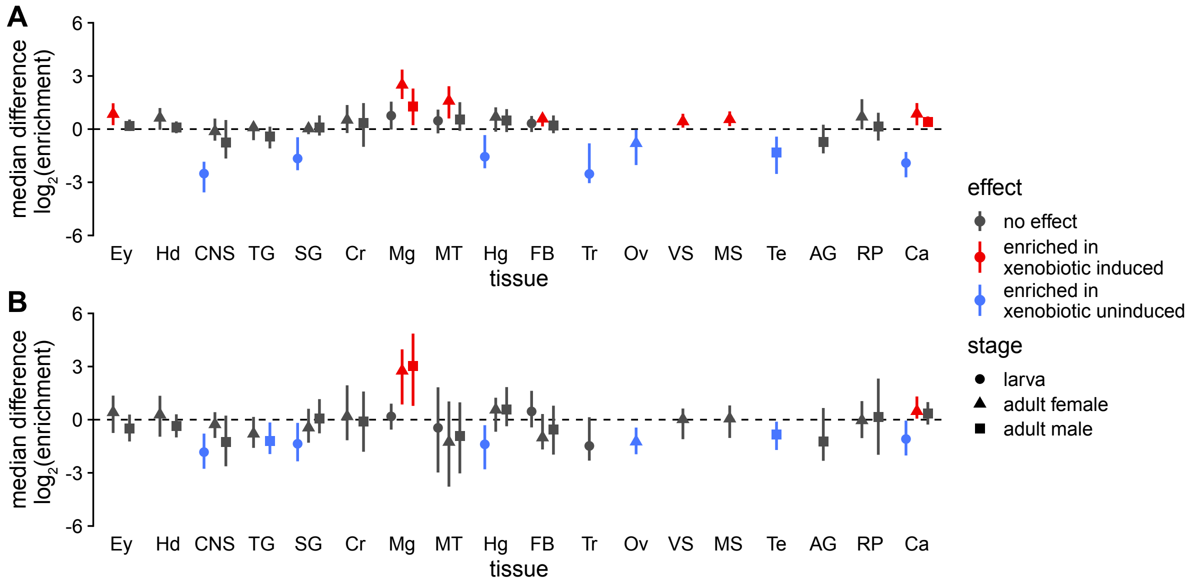


**Figure S6:** Median $\log_2$(enrichment) differences between xenobiotically induced and uninduced genes in the P450 (A) and EcKL (B) gene families for specific tissues and life stages. Error bars are 95% confidence intervals; effects were considered significant if the interval did not overlap with 0. Ey, eye; Hd, head; CNS, central nervous system; TG, Thoracicoabdominal ganglion; SG, salivary gland; Cr, crop; Mg, midgut; MT, Malpighian tubules; Hg, hindgut; FB, fat body; Tr, trachea; Ov, ovary; VS, virgin spermatheca; MS, mated spermatheca; Te, testis; AG, accessory gland; RP, rectal pad; Ca, carcass.

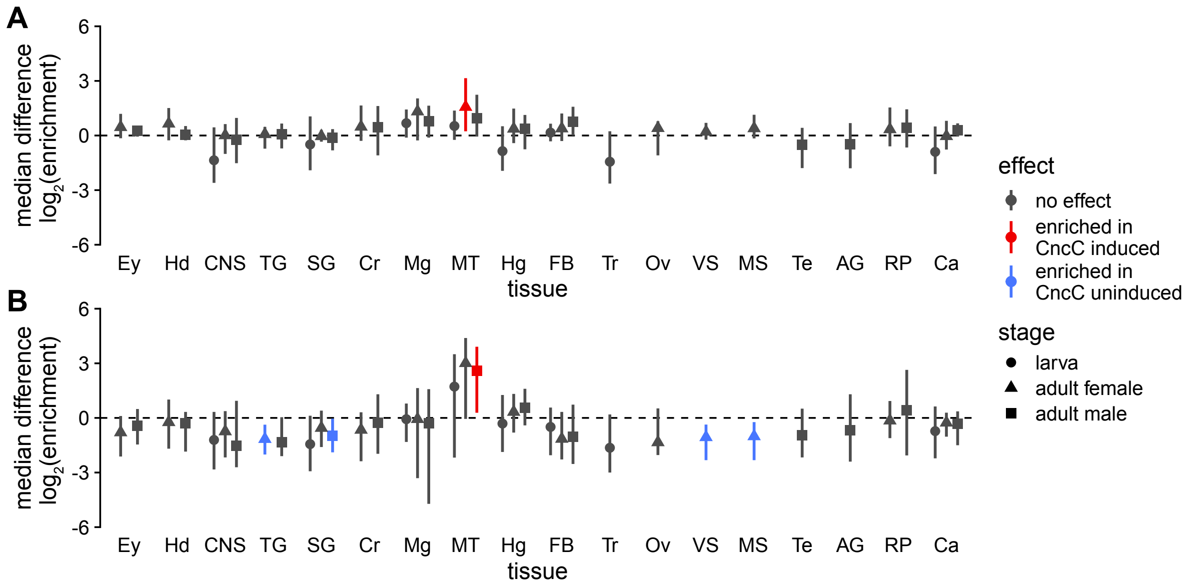


**Figure S7:** Median $log_2$(enrichment) differences between CncC induced and uninduced genes in the P450 (A) and EcKL (B) gene families for specific tissues and life stages. Error bars are 95% confidence intervals; effects were considered significant if the interval did not overlap with 0. Ey, eye; Hd, head; CNS, central nervous system; TG, Thoracicoabdominal ganglion; SG, salivary gland; Cr, crop; Mg, midgut; MT, Malpighian tubules; Hg, hindgut; FB, fat body; Tr, trachea; Ov, ovary; VS, virgin spermatheca; MS, mated spermatheca; Te, testis; AG, accessory gland; RP, rectal pad; Ca, carcass.

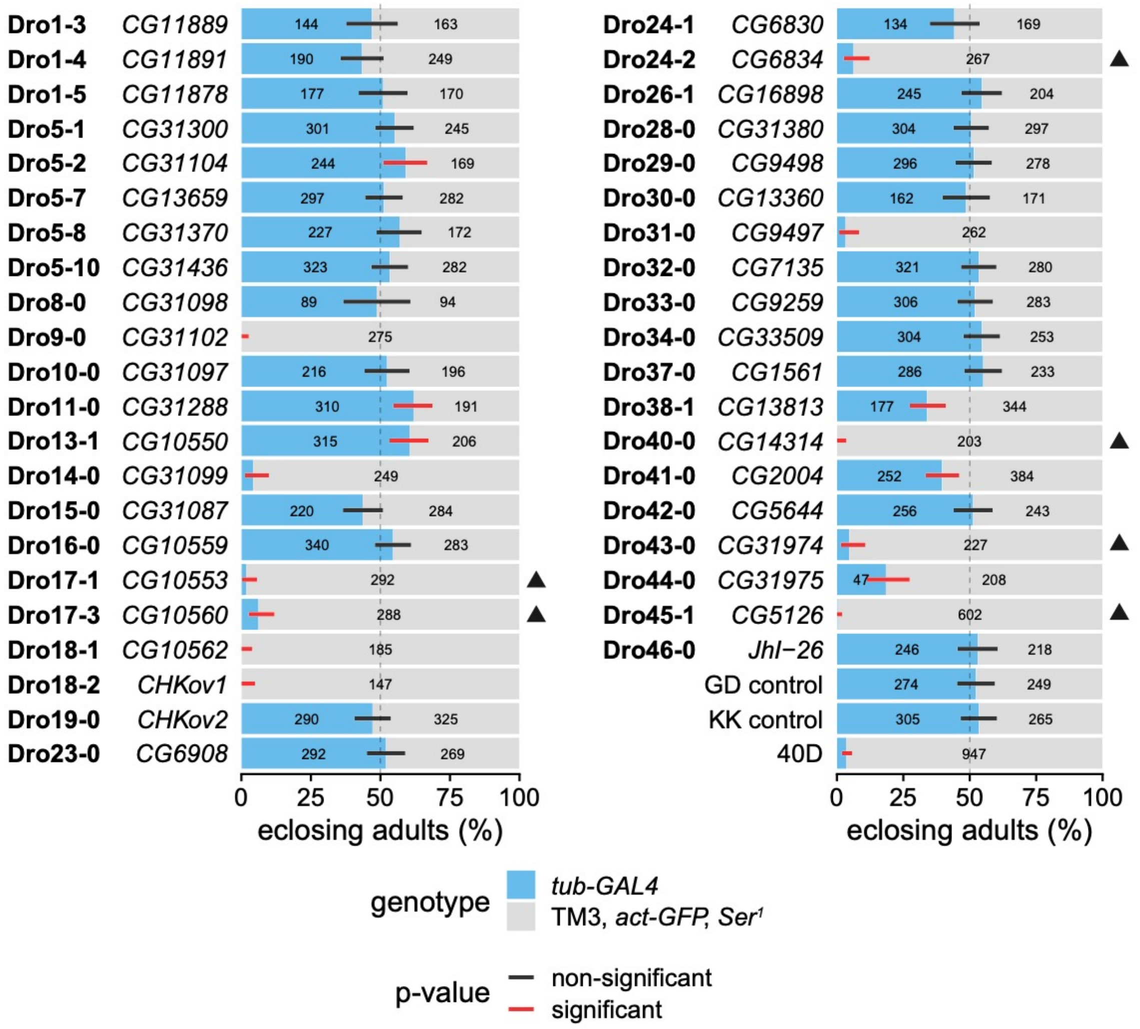


**Figure S8:** RNAi knockdown of 41 EcKL genes in *Drosophila melanogaster* and three control crosses, scoring offspring genotypes. Error bars are 99.89% confidence intervals (95% confidence interval adjusted for 44 tests) for the proportion of *tub-GAL4* individuals; black and red bars indicate non-significant or significant deviations, respectively, from expected genotypic ratios after correction for multiple tests. For the *CG31098* cross, the UAS-dsRNA construct was over a CyO balancer, so only non-CyO individuals have been scored. For the *CHKov1* cross, the UAS-dsRNA construct was on the X-chromosome, so only female individuals (inheriting the driver and responder) have been scored. The number of eclosed adults of each genotype is indicated by the number within each bar (numbers less than 40 are not shown). Results from KK library dsRNA lines with annotated insertions of the hairpin are indicated with black triangles; 40D is a line containing a UAS-only insertion at the annotated position for the KK library and acts as a positive control for *tiptop*-related phenotypes. dsRNA VDRC line IDs can be found in Table S4.
